# A sequence-specific RNA acetylation catalyst

**DOI:** 10.1101/2024.10.25.620255

**Authors:** Supuni Thalalla Gamage, Shereen Howpay Manage, Aldema Sas-Chen, Ronit Nir, Brett W. Burkhart, Isita Jhulki, Courtney N. Link, Manini S. Penikalapati, Jane E. Jones, Lakshminarayan M. Iyer, L. Aravind, Thomas J. Santangelo, Schraga Schwartz, Jordan L. Meier

## Abstract

N4-acetylcytidine (ac4C) is a ubiquitous RNA modification incorporated by cytidine acetyltransferase enzymes. Here, we report the biochemical characterization of *Thermococcus kodakarensis* Nat10 (TkNat10), an RNA acetyltransferase involved in archaeal thermotolerance. We demonstrate that TkNat10’s catalytic activity is critical for *T. kodakarensis* fitness at elevated temperatures. Unlike eukaryotic homologs, TkNat10 exhibits robust stand-alone activity, modifying diverse RNA substrates in a temperature, ATP, and acetyl-CoA-dependent manner. Transcriptome-wide analysis reveals TkNat10 preferentially modifies unstructured RNAs containing a 5’-CCG-3’ consensus sequence. Using a high-throughput mutagenesis approach, we define sequence and structural determinants of TkNat10 substrate recognition. We find TkNat10 can be engineered to use non-native acyl-CoA donors, providing insight into its cofactor specificity. Finally, we demonstrate TkNat10’s utility for site-specific acetylation of RNA oligonucleotides, enabling analysis of ac4C-dependent RNA-protein interactions. Our findings establish a framework for understanding archaeal RNA acetylation and a new tool for studying the functional consequences of ac4C in diverse RNA contexts.

## Introduction

Posttranscriptional modifications provide an evolutionary strategy to expand RNA function. One representative example is cytidine acetylation.^1^ In bacteria, archaea, and eukaryotes, the formation of N4-acetylcytidine (ac4C) within RNA is catalyzed by the TmcA, Kre33, and Nat10 family of cytidine acetyltransferase enzymes (Fig. 1). The architecture of these proteins is highly homologous across all three kingdoms of life and comprises at a minimum an N-terminal helicase domain, a Gcn5-related N-acetyltransferase (GNAT), and an RNA-binding domain all joined within a single polypeptide scaffold ranging from ∼600-1000 amino acids (Fig. 1).

**Figure 1.**
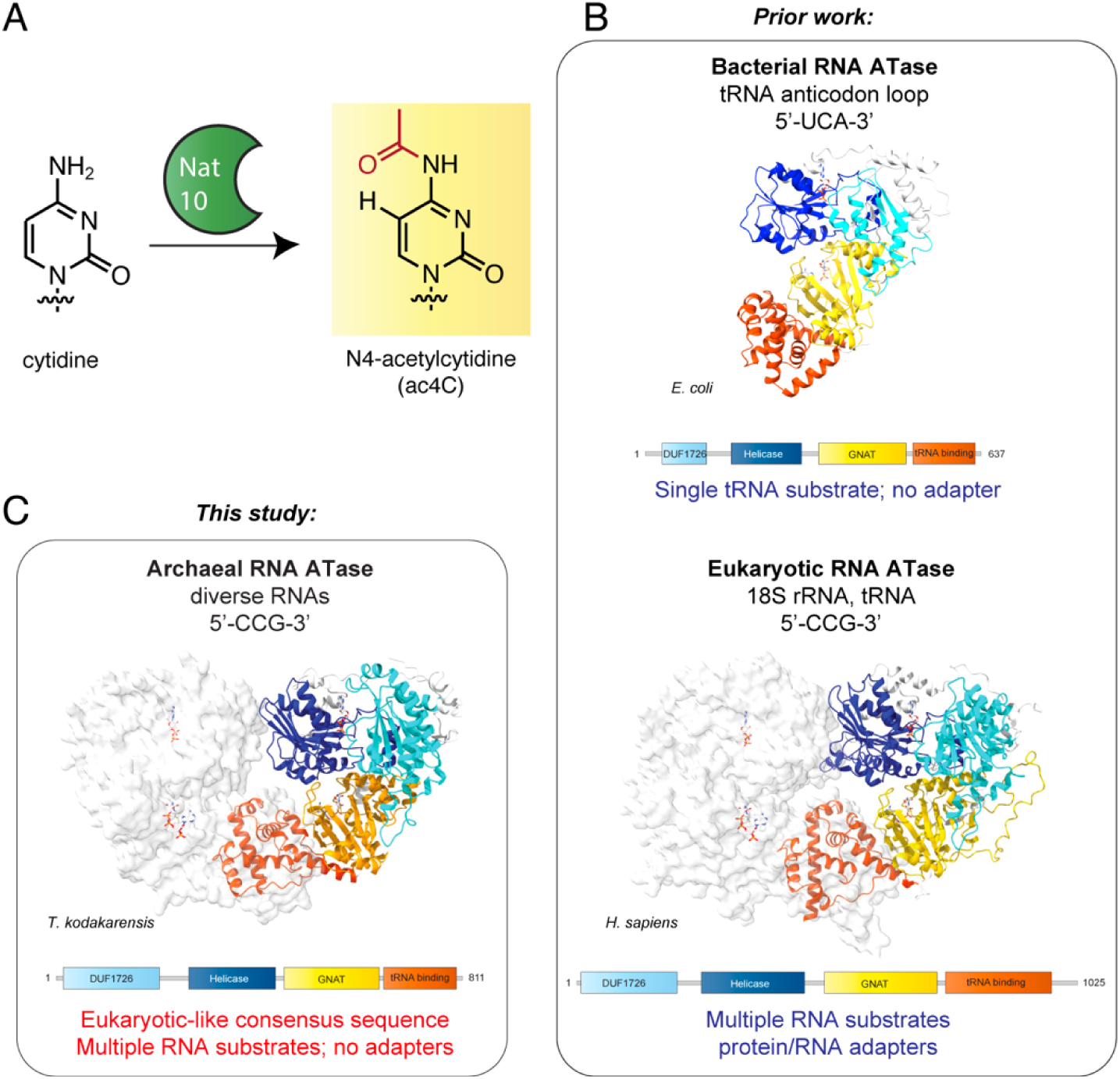
(A) RNA cytidine acetylation is catalyzed by the Nat10 family of enzymes. (B) Prior work has focused on characterization of bacterial RNA acetyltransferases that are highly specific or eukaryotic RNA acetyltransferases which use adapters to address 5’-CCG-3’ sequences in specific substrates. (C) This study characterizes an adapter-independent archaeal RNA acetyltransferase that modifies 5’-CCG-3’ consensus sequence in diverse substrates. *T. kodakarensis* Nat10 is depicted as a dimer based on AlphaFold Multimer predictions (Fig. S1).

In bacteria, ac4C helps ensure accurate protein translation through the modification of a single substrate, the wobble position of initiator tRNA^Met^.^2, 3^ Cytidine acetylation appears to also be highly controlled in eukaryotic organisms. Penetrant acetylation sites have thus far been validated within type II tRNA and 18S rRNA, where they have been implicated in tRNA stability and ribosome biogenesis, respectively.^4-6^ In contrast, archaea have demonstrated the potential for much more pervasive patterns of nucleic acid acetylation to exist. Applying nucleotide resolution ac4C sequencing as well as LC-MS to the archaeal hyperthermophiles *Thermococcus kodakarensis* and *Pyrococcus furiosis*, our group and others have detected hundreds of ac4C sites distributed across diverse coding and non-coding RNA substrates.^7-9^ In many substrates RNA acetylation was found to be directly proportional to growth temperature. This has led to the hypothesis that ac4C may aid fitness by increasing the robustness of RNA secondary structure at elevated temperature. Interestingly, in nearly all instances these sites occur at the central nucleobase of a 5’-C**<u>C</u>**G-3’ consensus sequence, the same motif that the human tRNA and rRNA sites are found within. How archaeal enzymes modify the same consensus sequence as eukaryotes – as well as the parameters of their vastly different substrate scope - remains to be established.

Despite much effort, biochemical studies of RNA cytidine acetyltransferase enzymes remain limited. *Escherichia coli* TmcA has been the most extensively characterized, revealing key determinants of tRNA^Met^ recognition, stand-alone ATPase activity, and a potential mechanism for communication between the helicase and GNAT active sites.^3,10^ Eukaryotic Nat10-type acetyltransferases require protein and short nucleolar RNA (snoRNA) adapters to efficiently modify their cellular tRNA and rRNA substates, and thus present a more challenging target for reconstitution.^11-13^ Our group and others have found sensitive radioactivity-based assays are necessary to detect the activity of reconstituted eukaryotic (fungal) Nat10.^5, 14, 15^ Although these studies have provided valuable biochemical and structural insights, their technical nature makes it difficult to analyze a large number of substrates, while the limited turnover of eukaryotic enzymes prevents an obstacle to preparative applications.^5^ Archaeal enzymes often provide tractable model systems useful for understanding more challenging eukaryotic counterparts as well as powerful tools for biotechnology.^16, 17^ However, the reconstitution and functional properties of RNA acetyltransferases involved in archaeal thermotolerance are currently unknown.

Here we report the biochemical characterization of the sequence-specific RNA acetylation catalyst *Thermococcus kodakarensis* Nat10 (TkNat10). This enzyme exhibits stand-alone RNA cytidine acetyltransferase activity and is able to modify substrates in a temperature, ATP, and acetyl-CoA-dependent fashion. Profiling the transcriptome-wide modification of *T. kodakarensis* substrates reveals the recombinant enzyme reaction directs acetylation to different substrates than are modified in vivo. A next-generation sequencing assay demonstrates a requirement for a 5’-CCG-3’ consensus sequence and a preference for unstructured substrates that can be influenced by secondary mutations. Finally, we probe the cofactor tolerance of this enzyme and demonstrate its promiscuous activity can be harnessed for site-specific acetylation of non-native RNA substrates, enabling us to study how ac4C perturbs RNA-protein interactions in its biologically relevant 5’-C**<u>C</u>**G-3’ context. Our studies define the properties of a novel component of the archaeal thermotolerance program and provide a tool for studying the functional consequences of cytidine acetylation in diverse RNAs.

## Material and Methods

### Expression and purification of recombinant TkNat10

The coding sequence of *Thermococcus kodakarensis* Nat10 (TK0754) was codon-optimized for *E. coli* expression, custom-synthesized (ATUM), and inserted into a bacterial expression vector with an N-terminal His and maltose binding protein (MBP) tag (His6-MBP-tev-TkNat10 WT; #24231-X02-566). The two TkNat10 mutants, His6-MBP-tev-TkNat10 S623G (#30594-X10-566) and His6-MBP-tev-TkNat10 S623G/W635A (#30594-X11-566), were similarly prepared. To prepare protein, each plasmid was transformed into BL21*pRARE competent E. coli cells (Protein Expression Laboratory, CCR/NCI). The proteins were expressed in cells grown at 37 °C until they were at an OD600nm of 0.6 and then induced with 500 μM IPTG, and grown overnight at 15 °C. After harvesting the cells by centrifugation for 30 min at 4 °C at 6800 g, they were lysed by sonication in a lysis buffer comprised of 50 mM Tris-HCl (pH 7.4), 150 mM sodium chloride, 10 mM Imidazole, 0.1 mg/mL lysozyme (Thermo Scientific), 125 U/mL Benzonase (Millipore), complete EDTA-free protease inhibitor cocktail tablets (Roche Diagnostics) and 2 mM TCEP. The lysate was pelleted in a pre-chilled centrifuge at 4 °C at 12000 rpm in a 14.5 rotor for 1 h. The resulting supernatant was filtered through a Millex HP 0.45 um 33 mm filter unit and, was loaded onto a HisTrap HP Ni column (Cytiva) pre-equilibrated with lysis buffer using a peristaltic pump. After washing with His binding buffer containing 50 mM Tris-HCl (pH 7.4), 150 mM sodium chloride and 25 mm imidazole, recombinant protein was eluted with a 10-250 mM gradient of imidazole in 50 mM Tris-HCl (pH 7.4), 150 mM sodium chloride. The collected fractions were further purified on a HiLoad 26/60 Superdex 200 column (GE Healthcare) pre-equilibrated with size exclusion chromatography (SEC) buffer consisting of 50 mM Tris HCl (pH 7.4), 300 mM NaCl, 10% glycerol, and 1 mm dithiothreitol. The purified protein was stored at −80 °C until use.

### Total RNA isolation from cells

Total RNA from human cells or *T. kodakarensis* cells (T559 strain, DTK0754:TkNat10 KO) was extracted using TRIzol (Invitrogen, cat. no. 15-596-018) according to the manufacturer’s protocol. Briefly, 1 ml TRIzol was used per 5–10×10^6^ mammalian or 1×10^7^ archaeal cells. After the cell lysis, samples were incubated at room temperature for 5 min followed by chloroform extraction and isopropanol precipitation. The RNA pellet was resuspended in water and quantified by UV absorbance and stored at −80°C.

### Biochemical assessment of TkNat10 activity

In vitro TkNat10 activity assays were done by mixing 50 mM Tris-HCl (pH 6), 400 mM NaCl, 5 mM MgCl_2_, 1 mM ATP, 0.2 mM Acetyl CoA, RNA substrate, ∼1 µM recombinant TkNat10 enzyme (∼ 10 µg), and water up to 100 µL. The reaction mixture was incubated at 65 °C for 3 h and the RNA was then isolated by using the Zymo RNA clean and concentrator-5 kit (Zymo Research, cat. no. R1013) according to the manufacturer’s protocol. For acetylation control, reactions were set up without the enzyme or acetyl CoA. Enzyme activity was also assessed as a function of pH and temperature using a wide variety of substrates, as described in the manuscript.

### Dot blot analysis

Dot blot assays were carried out to assess ac4C in RNA as described previously.^1^ RNA was purified after the TkNat10 reaction and spotted onto Amersham Hybond-N+ membranes (Cytiva, cat. no. RPN119B). Samples were crosslinked twice with 150 mJ/cm2 in the UV254nm Stratalinker 2400 (Stratagene, San Diego, CA). Following crosslinking, samples were blocked with 5% non-fat milk in TBST at room temperature for 30 min, and then probed with anti-ac4C (1:2000, Abcam) antibody in 5% non-fat milk in 1× TBST at 4 °C overnight. Membranes were washed three times with TBST and incubated with HRP-conjugated secondary anti-rabbit IgG (1:10,000 dilution, Cell Signaling) for 1 h. Membranes were washed four times with 1× TBST and developed with the SuperSignal ELISA Femto Maximum Sensitivity Substrate (ThermoScientific, cat. no. 37074) before detecting via chemiluminescent imaging.

### In vitro transcription (IVT) of TkNat10 substrates

In vitro transcription was performed with the HiScribe T7 Kit (New England Biolabs, cat. no. E2040S), according to the manufacturer’s instructions using DNA templates containing a T7 promoter upstream of the template sequence. Briefly, a 20 µL transcription reaction was set up with 100-200 ng DNA template, 10 mM of each NTP, 2 µL T7 RNA polymerase mix and 1× reaction Buffer. The reaction was incubated for 3□h at 37□°C, DNA was digested by adding 2 U of Turbo DNase (Invitrogen, cat. no. AM2238) directly to the IVT reaction and incubating for 15 min at 37□°C. Then the RNA was purified by using the Monarch RNA cleanup kit (NEB, cat. no. T2050L) according to the manufacturer’s protocol.

*In vitro transcription templates and products:*

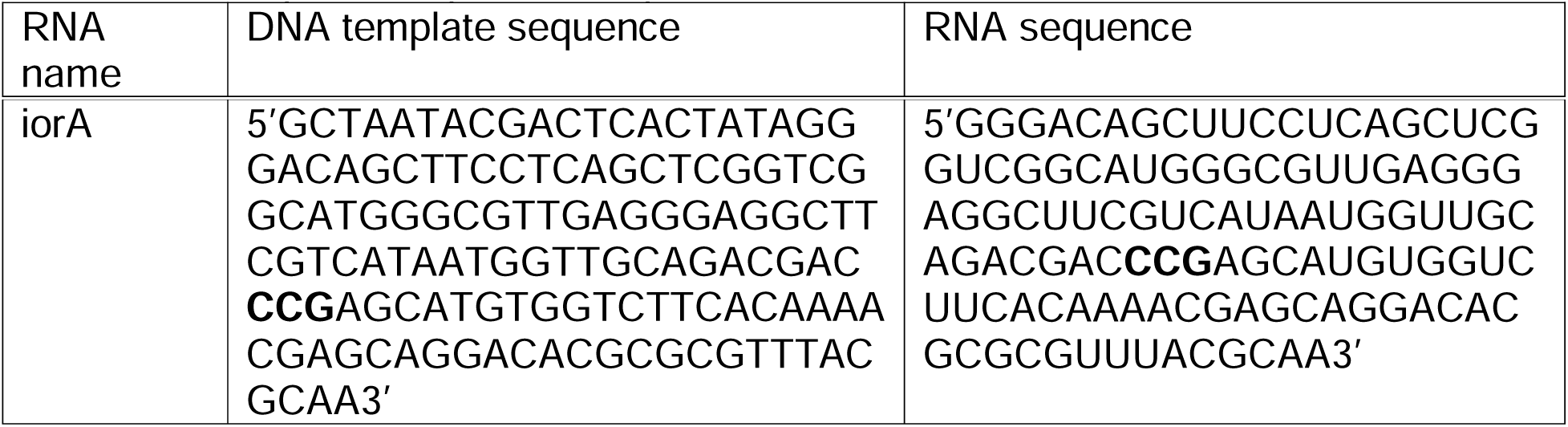

### Sanger sequencing-based analysis of TkNat10-catalyzed acetylation

Sanger-sequencing-based ac4C quantification was carried out as described previously.^2^ In vitro transcribed RNA was incubated with Turbo DNase (Invitrogen, cat. no. AM2238) for 30 min at 37 °C to remove any DNA contamination prior to the ac4C sequencing assay. Then, the iorA RNA (1 µg) was incubated with either sodium cyanoborohydride (100 mM in H_2_O) or vehicle (H_2_O) in a final reaction volume of 100 μL. Samples were incubated for 20 min at room temperature and quenched with 1 M Tris, pH 8 buffer (30 μL). Reactions were adjusted to 200 μL with H_2_O, ethanol precipitated, briefly dried on Speedvac, and resuspended in H_2_O. RNA from individual reactions (200 pg) was incubated with the iorA forward primer (ACAGCTTCCTCAGCTCGGTC, 0.2 uM) at 65 °C for 5 min and transferred to ice for 3 min to facilitate annealing in 1× Superscript III reaction buffer. After annealing, reverse transcription reactions were set up by adding 5 mM DTT, 200 units of Superscript III, optimized dNTP mix (250 µM GTP and 500 µM of ATP, CTP, TTP), 10 U of RNasin plus inhibitor (Promega), H_2_O up to 20 uL to the tubes and incubating for 60 min at 55 °C. PCR reactions using 2 μL of reverse transcription reactions were set up to amplify the cDNA product. To a 50 μL total reaction, 1× HF buffer, 0.5 μM of each forward and reverse primer (iorA fwd primer: ACAGCTTCCTCAGCTCGGTC, iorA rev primer: TTTGCGTAAACGCGCGTG), 200 μM of each dNTP, 2 U of Phusion hot start enzyme, and H_2_O were added. Thermocycling conditions: 30 sec at 98 °C, 33 cycles (10 sec at 98 °C, 30 sec at 65 °C, 20 sec at 72 °C) and 2 min extension at 72 °C. PCR products were run on a 2% agarose gel, stained with SYBR safe and visualized on UV transilluminator at 302 nm. Bands of the desired size were excised from the gel and DNA extracted using QIA-quick gel extraction kit from Qiagen and submitted for Sanger sequencing (Azenta) using the forward PCR primer. Processed sequencing traces were viewed using 4Peaks software. Peak height for each base was measured at the CCG and the percent misincorporation was determined using the equation: “Percent misincorporation= (T base peak intensity)/(sum of C and T base peaks)*100%” for sodium cyanoborohydride treated and vehicle reactions. The % misincorporation for each sample was calculated by subtracting the vehicle from sodium cyanoborohydride treated and then normalizing to empty vector and WT values.

### NGS-based analysis of TkNat10-catalyzed acetylation

To assess recombinant TkNat10’s ability to modify total RNA isolated from a *T. kodakarensis* TK0754 knockout strain, samples were treated with the enzyme (10 ug) followed by treatment with either NaCNBH_3_ or a mock-treatment (water/HCl) as described in the ac4C-seq protocol.^3^ Reverse transcription was performed using TGIRT at 42 °C for 16 hours.^4^ RNA was hydrolyzed and a second ligation performed adding a 3’ DNA oligo (5’ to the RNA strand) containing the Truseq R1 Illumina adapter and a 6bp UMI for BY4741/HEK293T/3T3 or without a UMI for mouse liver samples. Barcoded primers were then used to amplify the sequencing library via PCR. Libraries were subsequently sequenced on Illumina NovaSeq 6000 platform with an SP100 kit with read-lengths split evenly between R1 and R2. Read processing, alignment, and analysis were performed as previously described.^2^

### High-throughput mutational profiling of TkNat10 model substrates

DNA templates for the preparation of a degenerate library were purchased from Twist Bioscience (Table S4). Each template contained a T7 promoter at the 5’ end for in vitro transcription, along with either a human or *T. kodakarensis* h45 rRNA sequence, which was flanked by two stretches of DNA that did not contain any CCG motifs. The flanking DNA regions were added to increase RNA length for synthesis and sequencing purposes. The library contained h45 sequences with a single CCG motif (positions 4–6), which remained constant, while mutations were introduced into the surrounding h45 regions to generate single or double mutant substrates. Additionally, all single mutants that incorporated XCG or CCX sequences at the CCG motif were included in the library. Mutations that would result in the creation of additional CCG sites were excluded.

The DNA library was PCR-amplified using Phusion High-Fidelity (HF) DNA polymerase (annealing temperature = 56 °C, 33 cycles) with a fixed forward primer (5’-TAATACGACTCACTATAGGGTCGG-3’) and a reverse primer (5’-GACCTTTCCAAGGGCATAGATC-3’), which were common to all templates. Briefly, a 50 µL reaction was set up containing 1X HF buffer, 0.5 µM each of the forward and reverse primers, 200 µM of each dNTP, 2 U of Phusion Hot Start DNA polymerase, ∼17 ng of the template DNA pool, and nuclease-free water. The thermocycling conditions were as follows: initial denaturation at 98 °C for 30 seconds, followed by 33 cycles of 10 seconds at 98 °C, 30 seconds at 56 °C, and 20 seconds at 72 °C, with a final extension at 72 °C for 2 minutes. The amplified DNA library was run on a 2% agarose gel (expected product size: 170 bp), and the desired band was excised and purified using the NucleoSpin Gel and PCR Cleanup Kit (Macherey-Nagel, catalog no. 740609) following the manufacturer’s protocol.

In vitro transcription (IVT) was performed using the Superscript T7 Kit (New England Biolabs, catalog no. E2040S) according to the manufacturer’s instructions. In brief, a 100 µL transcription reaction was assembled using 1 µg of the DNA template pool, 10 mM of each NTP, 10 µL of T7 RNA polymerase mix, and 1× T7 reaction buffer. The reaction was incubated at 37 °C for 3 hours, after which 20 U of Turbo DNase (Invitrogen, catalog no. AM2238) were added to digest the DNA for 30 minutes at 37 °C. The IVT product was purified by denaturing it at 95 °C for 3 minutes in 1× RNA loading dye, followed by running the sample on a 6% PAGE-urea gel (14 W, 1× TBE). The RNA band (150 nucleotides) was visualized under UV light, excised, and subjected to a crush and soak method using 800 µL of buffer (500 mM ammonium acetate, 2 mM EDTA) at 4 °C overnight. The RNA-containing supernatant was recovered and desalted using a 1 kDa molecular weight cutoff (MWCO) centrifugal device (Pall Corporation, catalog no. MCP001C41).

The RNA library was then subjected to in vitro acetylation using *TkNat10*, as described previously. Each reaction used 3 µg of RNA and was conducted at 37, 55, 65, 75, and 85 °C for 3 hours. For the time-course experiments at 65 °C, reactions were incubated for 0.5, 1, 3, and 5 hours. Temperature-and time-dependent reactions were done in duplicate. Approximately 1 µg of RNA was used for CNBH_3_ treatment and mock (H_2_O treated) reactions (32 samples), followed by reverse transcription (RT), as described earlier. Superscript III-based RT reactions were performed with 8 ng of RNA (both CNBH_3_-treated and mock-treated) using the RT primer (5’-GACCTTTCCAAGGGCATAGATC-3’). The RT reactions were adjusted to a final volume of 25 µL with 5 µL of nuclease-free water, and the resulting cDNA was purified using MicroSpin G-25 columns (Cytiva, catalog no. 27532501) according to the manufacturer’s instructions.

The purified cDNA was then used in PCR to attach adapters for next-generation sequencing. PCR1 was performed using 5 µL of purified cDNA, 0.5 µM primers (forward primer: 5’-CCCTACACGACGCTCTTCCGATCTTCGGCCCACGGCCCTGGC-3’, reverse primer: 5’-GACTGGAGTTCAGACGTGTGCTCTTCCGATCGACCTTTCCAAGGGCATAGATC-3’), and Q5 Hot Start High-Fidelity 2× Master Mix (NEB, catalog no. M0494S) in a 50 µL reaction. The thermocycling conditions were as follows: initial denaturation at 98 °C for 30 seconds, followed by 5 cycles of 10 seconds at 98 °C, 30 seconds at 72 °C, and 20 seconds at 72 °C, with a final extension at 72 °C for 2 minutes. The amplified DNA was purified using PureLink PCR microspin columns (ThermoFisher, catalog no. K310050), eluted with 10 µL of nuclease-free water, and used as the template for PCR2.

In PCR2, i5 and i7 dual indexing primers were attached to the PCR1 amplicon using Q5 Hot Start High-Fidelity 2× Master Mix. The PCR2 forward primer was 5’-AATGATACGGCGACCACCGAGATCTACAC[i5]ACACTCTTTCCCTACACGACGCTCT TCCGATC*T-3’*, and the reverse primer was *5’-CAAGCAGAAGACGGCATACGAGAT[i7]GTGACTGGAGTTCAGACGTGTGCTCTTCC GATC*T-3’. Thermocycling conditions were 30 seconds at 98 °C, followed by 25 cycles of 10 seconds at 98 °C, 30 seconds at 72 °C, and 20 seconds at 72 °C, with a final extension at 72 °C for 2 minutes. The PCR products were purified using a 2% agarose gel (amplicon size: 283 bp), eluted with 15 µL of nuclease-free water, and diluted to 5 ng/µL (∼50 nM). The quality of the libraries was assessed using a High Sensitivity DNA Assay (Agilent, cat. no. 5067-4626) on an Agilent Bioanalyzer. Library size and concentration were obtained from the assay, and an equimolar concentration of the 28 libraries was pooled for sequencing. The sequencing-ready pool was quantified by qPCR before being subjected to a sequencing quality control (QC) run on an Illumina MiSeq sequencer. Data from the MiSeq run were analyzed, and adjustments to the libraries were made based on the read count for each sample. Following adjustment, the library pool was quantified again by qPCR, and a sequencing run was initiated on an Illumina NextSeq 2000 P2 paired-end 100-cycle sequencing platform. Basecalling was performed using Illumina RTA v3.10.30, and samples were demultiplexed using BCL2FASTQ v2.20. Quality control was performed using FastQC and FastQ_Screen, with all samples showing a Q30 base percentage greater than 75%. The samples yielded between 12 and 34 million pass-filter reads.

### Bioinformatic analysis of TkNat10 substrate library experiments

Sequences were uploaded and demultiplexed on the NCI Sequencing Facility’s database system with read quality assessed using FastQC. Target sequences were extracted from paired end sequencing data using the custom scripts R1_extract.perl (my $left_flank= "GTCGTAACAAGG"; my $max_length=24) and R2_extract.perl (my $left_flank= "CTCCCTTAATGATC"; my $right_flank = "CCTTGTTACGAC"). The R2 reads for each condition were transformed into their reverse complement (reverse_complement.perl), merged with the R1 reads for each condition (merge_files.perl) and the occurrence of each individual sequence in the merged reads for each condition counted (sequence_counts.perl; length = 24). The output of this analysis was then uploaded and further analyzed on the NIH HPC Biowulf Linux cluster. Counts for all 28 conditions were merged into a single parseable file (merge_counts.perl) and then filtered for matched occurrences of the 5’-CCG-3’ consensus sequence and 5’-CTG-3’ ac4C-seq sequence at positions 4-7. The percent CTG conversion was calculated by comparing the number of reads 5’-CTG-3’ at positions 4-7 to the total number of reads (5’-CCG-3’ + 5’-CTG-3’ for either i) NaCNBH_3_/HCl-treated versus vehicle-treated or ii) TkNat10-treated versus untreated samples for each condition. In practice the use of either control yielded nearly identical results and the +/- enzyme control was used. To correct for the fact that three mutants (U17 -> G17, U17 -> C17, and G20 -> C20) can create new TkNat10 consensus sequences, the percent CTG for these mutants was calculated combining the read counts for the converted and unconverted substrate sequences. Percent CCG remaining for each mutant was calculated analogously. Positional base frequencies were calculated from extracted 24-mers using Discriminative Regular Expression Motif Elicitation, part of the MEME Suite (https://meme-suite.org/meme/). Predicted free energies for library members were calculated using the ViennaRNA package (https://www.tbi.univie.ac.at/RNA/#) on the NIH HPC Biowulf Linux cluster. Scatterplots and heatmpas were plotted using Graphpad Prism. Custom perl scripts used in this study can be found at the GitHub repository, https://github.com/Meier-Lab-NCI/TkNat10.

### Identifying Modifications in RNA by MALDI Mass Spectrometry

To test the TkNat10 activity by MALDI mass spectrometry, a 10-mer RNA with the sequence CUUCCGUAGG (3136.9 Da) was purchased from IDT. In vitro TkNat10 reactions were carried out with either the WT TkNat10 enzyme, S623G or S623G/W635A variants in the presence of acetyl CoA or propionyl CoA cofactors using the activity assay described above. Control reactions were set up without CoA. After the reaction, the 10-mer RNA was purified using ziptip with C18 resin (Millipore, ZTC18S096) according to the manufacturer’s protocol. Samples for MALDI-MS were prepared by mixing 1-2 µL of RNA (∼100 ng) with 1-2 µL of 3-HPA matrix (prepared by mixing 9 parts 50 mg/mL 3-HPA in 50% acetonitrile/ddH O with 1 part 50 mg/mL ammonium citrate in ddH O). Samples were left to air dry on the MALDI plate, and mass spectra were recorded on a Shimadzu Biotech Axima Performance mass spectrometer in negative reflectron mode.

### RNA-protein pulldown

Both 3’-biotinylated and non-biotinylated 10-mer RNA oligonucleotides were purchased from Integrated DNA Technologies (IDT). These RNAs were acetylated using the TkNat10 enzyme, following previously described protocols. Control reactions without acetylation were set up by omitting acetyl-CoA from the enzymatic reaction. RNA purification was performed using ZipTips (Millipore, catalog no. ZTC18S096), according to the manufacturer’s instructions. Acetylation was confirmed via MALDI-TOF mass spectrometry.

For the pull-down experiments, whole-cell lysates were prepared from RAW 264.7 cells by resuspending cell pellets in 1x ice-cold PBS containing a protease inhibitor cocktail (EDTA-free, Cell Signaling Technology, cat. no. 5871). Cells were lysed by sonication using a QSonica Q500 sonicator (3-second pulse, amplitude 20%, with 30-second rests on ice between pulses, for a total of 5 pulses). Lysates were clarified by centrifugation at 20,817 g for 30 minutes at 4 °C and quantified using a Qubit 4.0 Fluorometer with the Qubit Protein Assay Kit (Thermo Fisher Scientific, cat. no. Q33211). The lysates were diluted to a final concentration of 3 mg/mL in lysis buffer and stored at -80 °C until further use. Prior to the pull-down assay, lysates were thawed on ice.

For each pull-down reaction, 500 µg of lysate was incubated with 10 µL of 1× protein-RNA binding buffer, 30 µL of 50% glycerol, 100 pmol of yeast tRNA^Phe^, 0.8 U/µL of RNasin, and nuclease-free water to a final volume of 100 µL. The RNA-protein pull-down assay was performed using the Pierce Magnetic RNA-Protein Pull-Down Kit, following the manufacturer’s protocol. Magnetic streptavidin beads were first washed twice with 20 mM Tris-HCl (pH 7.5). Subsequently, 50 µL of 1× RNA capture buffer (20 mM Tris-HCl, pH 7.5, 1 M NaCl, 1 mM EDTA) and 0.8 U/µL of RNasin were added to the beads, followed by incubation at room temperature for 30 minutes. After removing the supernatant, the beads were pre-blocked by incubating with 50 pmol of yeast tRNA^Phe^ at 4 °C for ∼16 hours while rotating.

After pre-blocking, the supernatant was discarded, and 50 pmol of biotinylated RNA in 1× RNA capture buffer was added to the beads, gently mixed, and incubated for 30 minutes at room temperature with agitation. Following RNA binding, the beads were washed twice with 20 mM Tris-HCl (pH 7.5) and once with 1× protein-RNA binding buffer (0.2 M Tris-HCl, pH 7.5, 0.5 M NaCl, 20 mM MgCl□, 1% Tween-20). The beads were then incubated with the prepared cell lysate solution for 1 hour at 4 °C. Control reactions were performed by incubating lysates with beads that had no biotinylated RNA bound. After protein capture, the beads were washed three times with wash buffer (20 mM Tris-HCl, pH 7.5, 10 mM NaCl, 1% Tween-20) and then exchanged into 50 mM HEPES-KOH (pH 7.5) for subsequent proteomic analysis.

### LC/MS/MS and Proteomic analysis of the RNA-protein pulldown samples

#### Digestion of biot-10mer on-bead proteins and TMTpro labeling

Each sample bound to capture beads in 50 µL of 50 mM HEPES pH 8 was treated with 150 μL of digestion buffer containing the following composition: 10 mM TCEP, 40 mM chloroacetamide, and 15 ng/µL trypsin/LysC in the lysis buffer provided with the EasyPep kit (Thermo, cat. No. A40006). Samples were incubated at 37 °C, overnight in the dark shaking at 1000 rpm. After digestion 150 μL of each solution was transferred to a new tube and treated with 10 µL of 10 μg/μL TMTpro (Thermo, cat. no. A52045) reagent and incubated for 1 h at 25 C with shaking. Excess TMTpro was quenched with 50 μL of 5% hydroxylamine, 20% Formic acid for 10 min and samples were combined and cleaned using EasyPep mini columns (Thermo, cat. No. A40006) according to the manufacturer’s protocol. Eluted peptides were dried in speed-vac.

#### LC/MS analysis of peptides

Dried peptides were resuspended in 50 μL of 0.1% FA and 5 μL was analyzed in duplicate using a Dionex U3000 RSLC in front of a Orbitrap Eclipse (Thermo) equipped with an EasySpray ion source. Solvent A consisted of 0.1% FA in water and Solvent B consisted of 0.1% FA in 80% ACN. The loading pump consisted of Solvent A and was operated at 7 μL/min for the first 6 minutes of the run then dropped to 2 μL/min when the valve was switched to bring the trap column (Acclaim™ PepMap™ 100 C18 HPLC Column, 3 μm, 75 μm I.D., 2 cm, PN 164535) in-line with the analytical column EasySpray C18 HPLC Column, 2 μm, 75 μm I.D., 25 cm, PN ES902). The gradient pump was operated at a flow rate of 300 nL/min and each run used a linear LC gradient of 5-7% B for 1 min, 7-30% B for 83 min, 30-50% B for 25 min, 50-95% B for 4 min, holding at 95% B for 7 min, then re-equilibration of analytical column at 5% B for 17 min. All MS injections employed the TopSpeed method with three FAIMS compensation voltages (CVs) and a 1 second cycle time for each CV (3 second cycle time total) that consisted of the following: Spray voltage was 2200 V and ion transfer temperature of 300 C. MS1 scans were acquired in the Orbitrap with resolution of 120,000, AGC of 4e5 ions, and max injection time of 50 ms, mass range of 350-1600 m/z; MS2 scans were acquired in the Orbitrap using TurboTMT method with resolution of 15,000, AGC of 1.25e5, max injection time of 22 ms, HCD energy of 38%, isolation width of 0.4 Da, intensity threshold of 2.5e4 and charges 2-6 for MS2 selection. Advanced Peak Determination, Monoisotopic Precursor selection (MIPS), and EASY-IC for internal calibration were enabled and dynamic exclusion was set to a count of 1 for 15 sec. The only difference in the methods was the CVs used, one method used CVs of -45, -60, -75 and the second used CVs of -50, -65, -80.

#### Database search and post-processing analysis

Both injections were batched together as fractions and searched with Proteome Discoverer 2.4 using the Sequest node. Data was searched against the Uniprot Human database from Feb 2020 using a full tryptic digest, 2 max missed cleavages, minimum peptide length of 6 amino acids and maximum peptide length of 40 amino acids, an MS1 mass tolerance of 10 ppm, MS2 mass tolerance of 0.02 Da, variable oxidation on methionine (+15.995 Da) and fixed modifications of carbamidomethyl on cysteine (+57.021), TMTpro (+304.207) on lysine and peptide N-terminus Percolator was used for FDR analysis and TMTpro reporter ions were quantified using the Reporter Ion Quantifier node and normalized on total peptide intensity of each channel. TMTpro channel assignment for conditions are as follows: 1) Control-biot-10mer: 126, 127N, 128N; 2) ac4c-biot-10mer: 129N, 130N, 131N; 3) Lysate: 132N, 133N, 134N.

### Biochemical reconstitution of an archaeal RNA acetyltransferase

As a preface to *T. kodakarensis* Nat10 reconstitution, we first sought to determine the necessity for its catalytic activity in vivo. Prior research has shown that genetic deletion of TkNat10 causes decreased fitness at elevated temperatures.^7, 8^ One caveat to this finding is that TkNat10 contains multiple protein domains (Fig. 1), making it unclear whether this phenotype was specifically due to loss of acetyltransferase activity. To address this, we engineered a *T. kodakarensis* strain in which the highly conserved histidine (H506) of the ‘HY’ motif in the GNAT domain was mutated to alanine (Fig. 2A). This motif, present in all characterized Nat10-type GNAT domains, is essential for the activity of bacterial and eukaryotic enzymes.^6, 10, 15^ Analysis of ac4C content in total RNA isolated from the H506A strain confirmed complete loss of acetylation (Fig. S2A). Monitoring growth as a function of time revealed that strains lacking RNA acetylation – either due to catalytic mutation or genetic deletion – exhibit significantly decreased growth at 85 °C and 95 °C (Fig. 1A-B, Fig. S2B-C). These findings validate the critical role of TkNat10’s cytidine acetyltransferase activity in *T. kodakarensis* fitness under thermal stress.

**Figure 2.**
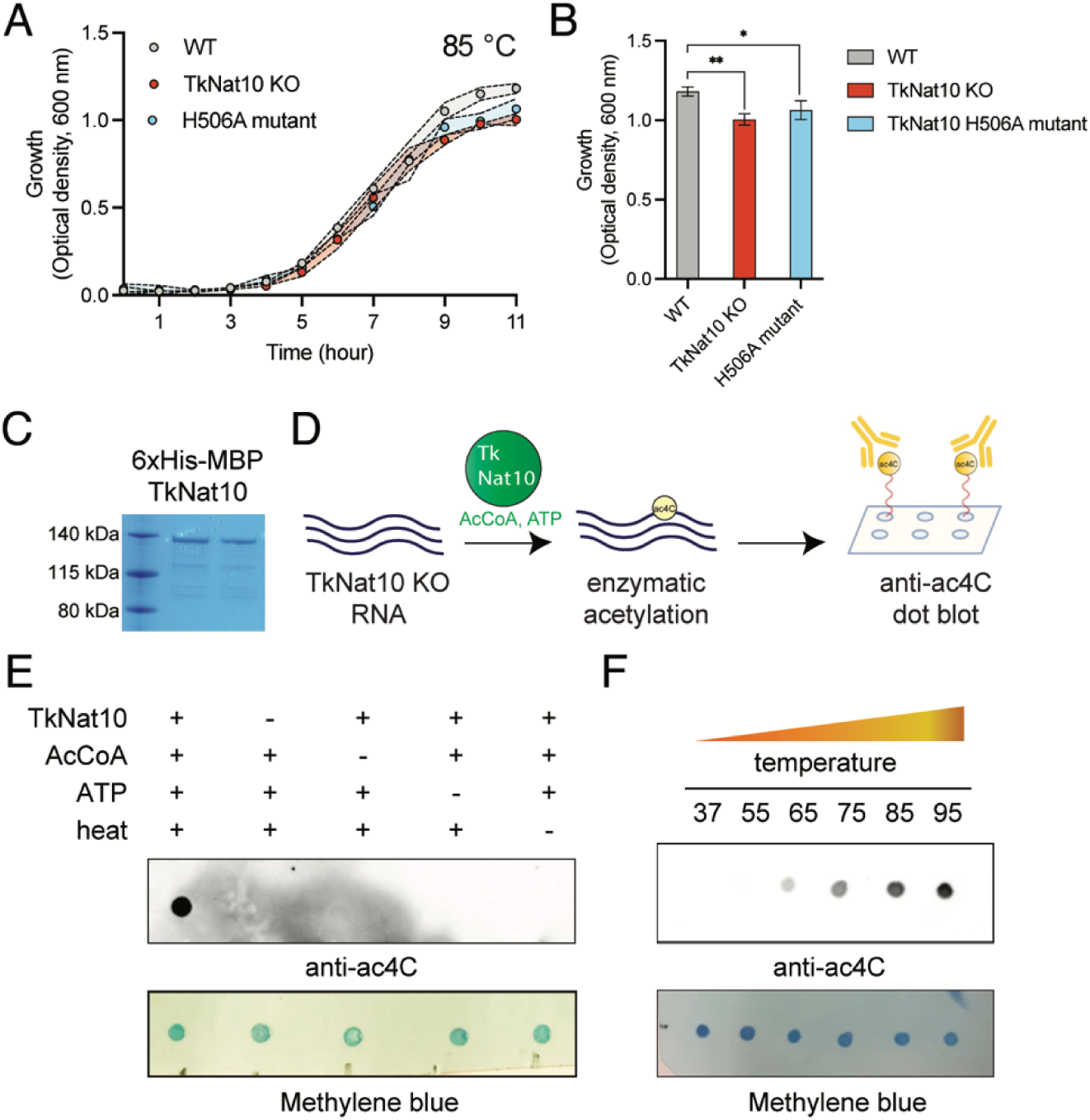
(A) Growth rate of WT *T. kodakarensis*, TkNat10 deletion strain, and strain containing a TkNat10 H506A catalytic mutant at 85 °C. (B) Differential growth of *T. kodakarensis* strains at 85 °C for 11 hours. Values represent ≥ 3 replicates, analyzed by two-tailed student’s t-test (*P<0.05, **P<0.01). (C) Purification of 6xHis-tagged TkNat10 enzyme. (D) Schematic for qualitative analysis of TkNat10 activity by anti-ac4C dot blot. (E) Acetylation-catalyzed by TkNat10 is enzyme, acetyl-CoA, ATP, and heat-dependent. (F) TkNat10 catalysis as a function of temperature.

Having established the in vivo importance of TkNat10, we proceeded to its in vitro reconstitution. An *E. coli* codon-optimized version of the TkNat10 gene (TK0754) was synthesized and inserted into a Gateway cloning vector and used to produce N-terminal His and His/MBP tagged versions of the enzyme via bacterial overexpression. Soluble protein was purified by immobilized metal chromatography and FPLC. The His and His/MBP versions displayed roughly equivalent activity (vide infra); for practicality we will describe the characterization of His/MBP-tagged TkNat10, which consistently gave higher yields of soluble protein. The purified 1207 amino acid enzyme showed a molecular weight of ∼137 kDa on sodium dodecyl sulfate polyacrylamide gel electrophoresis (SDS PAGE), aligning with its predicted mass (Fig. 2C, Fig. S3).

TkNat10 is known to modify hundreds of RNA targets in vivo,^1^ which complicates the selection of an optimal biochemical substrate. To circumvent this, we developed a non-radioactive assay for RNA acetyltransferase activity using total RNA isolated from *T. kodakarensis* strains deficient in TkNat10 (TkNat10 KO). Our rationale was that RNA from these strains, being entirely deficient in ac4C,^7^ should contain many substrates amenable to TkNat10 modification that could be readily detected by anti-ac4C dot blot (Fig. 2D).^18, 19^ Incubation of recombinant TkNat10, acetyl-CoA (AcCoA), and ATP with KO RNA at 65°C produced a robust ac4C signal (Fig. 2E). Omission of any single reaction component (enzyme, AcCoA, ATP) or heat abolished the signal. TkNat10-catalyzed acetylation was temperature-dependent, with RNA modification increasing over a temperature gradient of 55 °C to 95 °C (Fig. 2F). RNA acetylation was further found to be installed in a time-dependent manner with an optimal pH of 6 -7 (Fig. S4). These studies establish a strategy for heterologous expression and biochemical reconstitution of archaeal RNA acetyltransferase activity.

### Transcriptome-wide analysis of recombinant RNA acetyltransferase activity

To gain a higher resolution picture of the activity of recombinant TkNat10 we next analyzed the biochemically modified pool using quantitative ac4C sequencing (ac4C-seq; Fig 3A).^20^ This approach exploits the unique electronic character of ac4C which renders it susceptible to acid-catalyzed sodium cyanoborohydride reduction. The product of this reaction – reduced ac4C – causes misincorporations during reverse transcription and produces a strong mutational signature at ac4C sites that can be detected upon cDNA sequencing.^21^ Consistent with dot blot data, applying ac4C-seq to TkNat10 KO RNA treated with enzyme and cofactors at 75 °C revealed the formation of ac4C misincorporation signals (>5%) across 598 RNA sites (Table S1). In line with the known cellular specificity of this enzyme, these signals were observed exclusively within 5’-C**<u>C</u>**G-3’ sequences. To systematically compare the modification profile of biochemical and cellular TkNat10, we took advantage compare the modification profile of biochemical and cellular TkNat10, we took advantage of the quantitative nature of ac4C-seq which allows misincorporation percentage to be used as a proxy for modification level. We identified 236 ac4C sites within the *T. kodakarensis* transcriptome that were quantified (depth ≥50, misincorporation > 0) both in the in vivo experiment^7^ as well as upon enzymatic TkNat10 treatment. In vivo many of the most penetrant ac4C sites lie within rRNA and tRNA (Fig. 3B, y-axis). In contrast, the highest levels of misincorporation driven by recombinant TkNat10 treatment were found in messenger RNA (mRNA; Fig. 3B, x-axis). To validate this finding, we in vitro transcribed a highly modified mRNA encoding indolepyruvate ferredoxin oxidoreductase, alpha subunit (iorA; Fig. 3C). Consistent with our transcriptome-wide analysis, we found that treatment with recombinant TkNat10 catalyzed significant modification of this in vitro transcription-produced substrate (Fig. 3C). Interestingly, secondary structure prediction indicates the modification site in iorA occurs in an unpaired 5’-CCG-3’ motif adjacent to two stem loops. As coding transcripts are often less structured than non-coding RNAs, we next asked: what is the relationship between folding energy and TkNat10 modification? To assess this, we divided the 236 RNAs modified by TkNat10 into five bins based on their degree of misincorporation, ranging from high to low. The Vienna RNA package was then used to calculate the minimum free energy (MFE) of a 100-nucleotide fragment of each RNA centered on the modified 5’-CCG-3’ site.^22^ RNAs exhibiting the highest modification by TkNat10 (e.g. bin 1, Fig. 3D) on average had the least negative MFE, suggestive of less structure. These data indicate the wide substrate scope of recombinant TkNat10 and suggest its stand-alone activity may be anticorrelated with RNA secondary structure.

**Figure 3.**
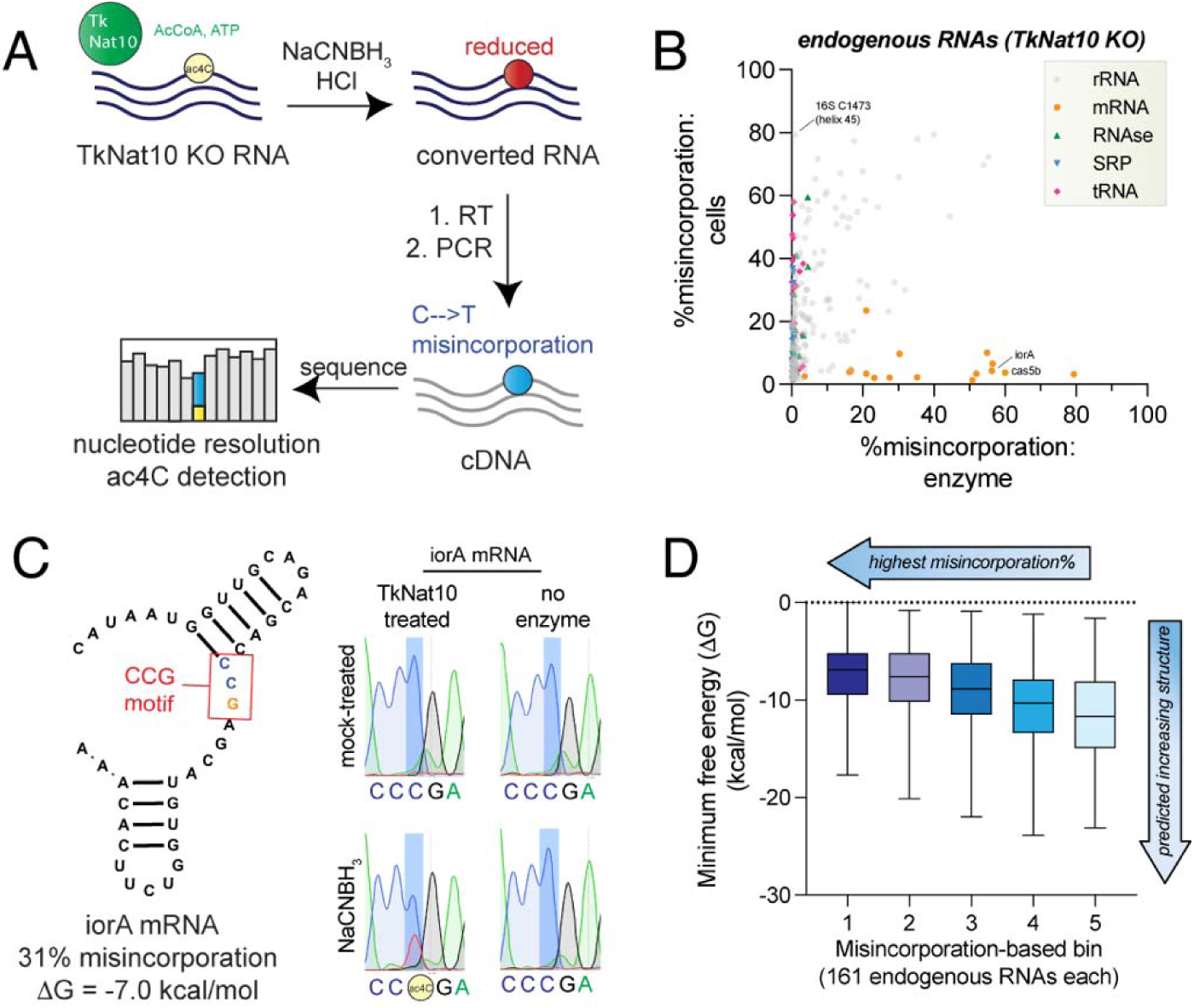
(A) Schematic for transcriptome-wide identification of recombinant TkNat10 substrates in total RNA isolated from a TK0754 knockout (KO) strain. (b) Correlation of misincorporation percentages observed upon treatment of KO RNA with recombinant TkNat10 enzyme (x-axis) or in RNA isolated from wild-type *T. kodakarensis* (y-axis). (C) Example of an enzyme-catalyzed misincorporation site in iorA mRNA which was validated by amplicon-based Sanger sequencing. (D) Correlation of ac4C-seq misincorporation percentage with minimum free energy (MFE) of the 30 base region surrounding the ac4C site calculated by ViennaRNA.

### High-throughput mutagenesis of a TkNat10 substrate

While our transcriptome-wide experiment describes the ability of recombinant TkNat10 to modify a wide variety of *T. kodakarensis* RNAs, it does not provide a detailed view into how the sequence of any individual substrate impacts acetylation. To evaluate this in a systematic fashion, we developed a strategy to apply ac4C-seq for high-throughput mutational profiling of model substrates. The approach harnesses advances in parallel oligonucleotide synthesis to construct large libraries of substrate sequences which can be in vitro transcribed, treated with TkNat10 and cofactors, and subjected to ac4C-seq. As initial targets for analysis, we focused on the highly conserved ac4C sites found in helix 45 of *T. kodakarensis* and *H. sapiens* small subunit rRNA (Fig. 4A). Our hypothesis was that these two models would enable analysis of evolutionary conservation and divergence of substrate recognition in a well-defined structural context. Furthermore, the presence of two non-canonical G • U base pairs in the stem of the human substrate would allow us to test the effect of strengthening base pairs on TkNat10 activity.

**Figure 4.**
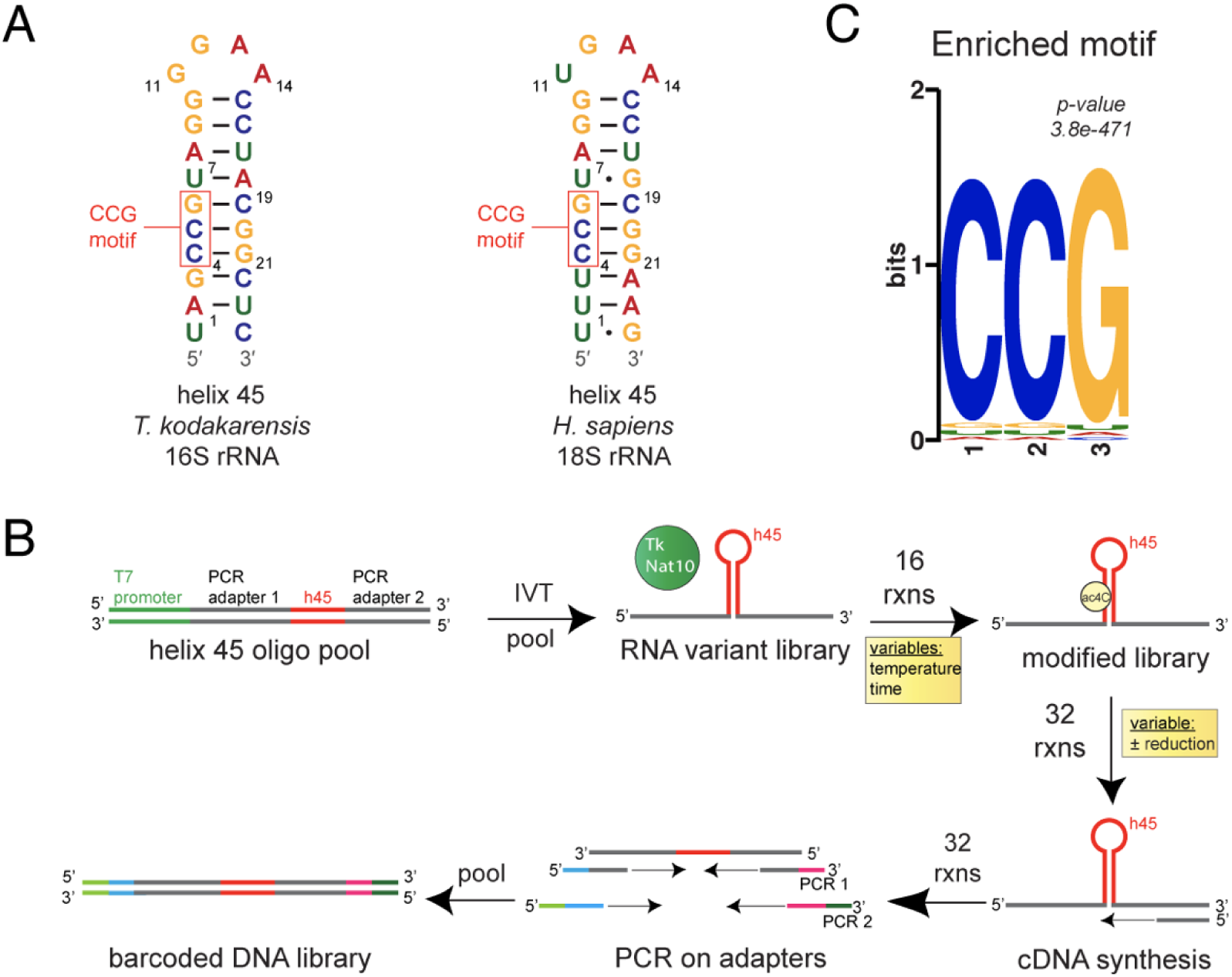
(A) Structure of 16S and 18S SSU helix 45 rRNA model substrates. (B) Schematic for high-throughput analysis of TkNat10 substrate-specificity using a hairpin substrate. (C) STREME analysis of 3-bp motifs differentially enriched in reaction (NaCNBH_3_-treated) and control (mock-treated) libraries. Both libraries were subjected to equivalent ac4C sequencing reactions.

Accordingly, we first designed 24-mer sequences which kept the the 5’-CCG-3’ found at positions 4-6 constant (numbering from the base of the helix) while sampling all single mutations for both models. Mutations that would create potential 5’-CCG-3’ substrate sequences were omitted to simplify analysis. To enhance our analysis of human helix 45, we additionally surveyed all possible pair-wise mutants as well and positions 4/6, with the aim of identifying epistatic mutations and probing TkNat10’s necessity for the 5’-CCG-3’ consensus, respectively. T7 templates encoding the designed rRNA hairpin as well as PCR adapters were purchased, PCR-amplified, subjected to in vitro transcription. Purified RNA was incubated with recombinant TkNat10 and cofactors and subjected to ac4C-seq (Fig. 4B). In addition to a ‘no reduction’ control, we examined the effect of time (0.5-5 h) and temperature (37-85 °C) on conversion. Analyzing a control library where no conversion was expected (no reduction, 37 °C) revealed adequate coverage (>200 reads) of all single mutants of the parent sequence, with a median depth of 2264 reads per library member for this single condition (Table S2).

To extract TkNat10 activity from our sequencing data reads containing 5’-CCG-3’ and 5’-CTG-3’ at bases 4-6 were paired and used to calculate ‘% CCG conversion’ (Table S2). This ratiometric measure normalizes for variable read depth caused by the differential representation of each sequence in the initial library. Our expectation was that the most efficient TkNat10 substrates would display a high ‘% CCG conversion’. To correct our data for heat-catalyzed deamination, another potential contributor to conversion, we subtracted the conversion rate of samples treated with TkNat10/cofactors/heat but <u>not</u> reduced. Simple, Thorough, Rapid, Enriched Motif Elicitation (STREME)^23^ analysis was used to identify 3-mers overrepresented in non-reduced versus sodium cyanoborohydride-reduced samples and readily identified the 5’-CCG-3’ acetylation consensus sequence, consistent with the expected activity of the enzyme (Fig. 4C).^24^

Comparing activity of TkNat10 towards the parent *T. kodakarensis* and *H. sapiens* SSU rRNA sequence (Fig. 5A and Fig. 5B, ‘0’ left-most column) some immediate trends emerged. Both substrates show temperature-dependent modification that is driven to near completion after 3 h of incubation at 85 °C. The human helix 45 surrogate displays time-dependent modification at 65 °C, while the more GC-rich *T. kodakarensis* hairpin is significantly modified at 75 °C and not lower temperatures. None of the mutations enabled modification at 37 °C. Scanning across the 5’ hairpin stem, we observe two mutations, U1 -> G1 in *T. kodakarensis* and U1 -> C1 in *H. sapiens* that clearly decrease ac4C formation (Fig. 5C). U7 -> C7 in *H. sapiens* shows an even more dominant effect, limiting ac4C deposition even at 75 °C. Each of these mutations install a more stable G • C pair within the helix. This indicates stabilized duplex hairpin structures are less susceptible to TkNat10-catalyzed modification. A similar, albeit less consistent, effect can be seen upon mutation of *H. sapiens* G18 -> A and G24 - > A, which replace the G • U with a canonical A • U pair. All other mutations in our library targeted to the stem disrupt canonical base-pairing and largely had neutral or positive effects on acetylation. Common stem mutations which improved modification of both the *T. kodakarensis* and *H. sapiens* substrates were observed at positions 16, 17, 19, 20, and 21 in the 3’ stem (Fig. 5A-B). The latter three (19-21) are positioned anti-parallel to the 5’-CCG-3’ consensus sequence (‘anti-CCG’) and appeared among the most conducive to modification (Fig. 5C, right). TkNat10’s ability to modify an unpaired substrate cytidine stands in marked contrast to the recently characterized Kre33/Tan1 complex, which shows an absolute requirement for base-pairing at the modified base.^14^ Tetraloop mutations influenced acetylation to a relatively minor extent, with the highest ranking sequences containing purine pairs at the internal position (Fig. S5).

**Figure 5.**
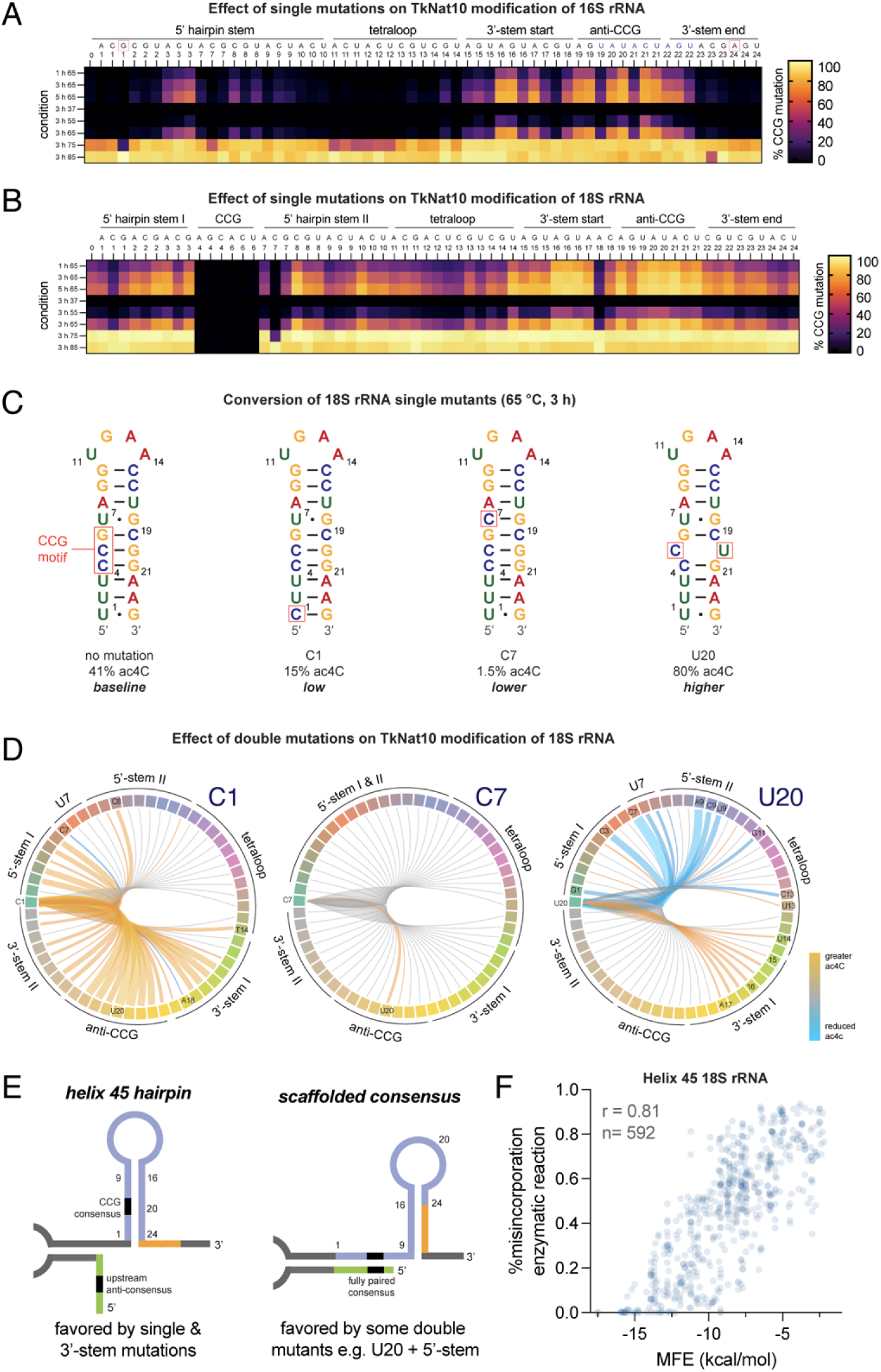
(A) Heatmap depicting % misincorporation (‘% CCG mutation’) as a function of time and temperature for 16S rRNA substrate. Each column corresponds to a mutant sequence with the mutation indicated in the top row of the header and the position in the bottom row of the header. Data represents the average of two replicate experiments. Raw data is given in Table S2. (B) Heatmap depicting % misincorporation (‘% CCG mutation’) as a function of time and temperature for 18S rRNA substrate. Each column corresponds to a mutant sequence with the mutation indicated in the top row of the header and the position in the bottom row of the header. Data represents the average of two replicate experiments. Raw data is given in Table S2. (C) Predicted secondary structure of 18S SSU helix 45 rRNA substate as well as C1, C7, and U20 mutants. (D) Circular arc plot depicting interaction of secondary mutations with C1, C7, and U20 mutations. Interaction scores were calculated from difference in summed % CCG conversion for parent and double mutant across three conditions (3 h 55 °C, 3 h 65 °C, and 5 h 65 °C). The width and intensity of each arc is scaled to the strength of the interaction, with mutations that stimulate ac4C modification in orange and those that reduce it in blue. Secondary mutations with a small effect size (averaging 10% or less) were depicted in grey. (E) Simplified schematic indicating the potential for 18S helix 45 mutation to form alternative RNA structures by interacting with scaffolding RNA. Left: The expected helix 45 surrogate forms and is predictably modulated by single mutations and double mutants in the 3’ stem. Right: When weakened helix 45 variants (e.g. U20) acquire a second mutation in the 5’ stem – particularly at position 9 – they favor an alternative structure in which the consensus sequence becomes fully paired with an upstream sequence, reducing susceptibility to TkNat10 modification. Detailed exemplary structures provided in Fig. S7. (F) Correlation of ac4C-seq misincorporation percentage with minimum free energy (MFE) for 18S rRNA variants. To focus exclusively on the helix 45 hairpin this analysis was tailored to mutants <u>not predicted</u> to form the alternative structure and included i) all single mutants, ii) mutants that strengthen the helix by forming new base pairs (e.g. C1, A18), and iii) position 15-16 or 19-21 mutants with a secondary mutation in stem II. The misincorporation is derived from incubation of TkNat10 and cofactors with RNA at 3 h 65 °C. C7 mutants are uniformly not modified under these conditions and thus were omitted from the graph for clarity.

Our inclusion of all double mutants in our *H. sapiens* helix 45 library provides the opportunity to understand the interplay of individual variants. We began by examining the two variants that form new C • G base pairs, U1 -> C1 and U7 -> C7. To evaluate the effect of secondary mutations on modification rate, we calculated the difference in summed % CCG conversion for parent and double mutant across three conditions (3 h 55 °C, 3 h 65 °C, and 5 h 65 °C). As conversion was incomplete under each of these conditions, we hypothesized we could use this as a crude measure of relative enzyme modification rate. These values were graphed as arc plots, with double mutants showing increased modification rate relative to parent in orange, those similar in effect in grey, and those showing decreased modification rate relative to parent in blue (Fig. 5D). The decreased modification caused by C1 mutation is rescued by adjacent mutations at positions 2 and 3 as well as a wide variety of 3’-stem variants, most notably at position 16, 20, and 21 (Fig. 5D, left, Fig. S6A). In contrast, the C7 mutation was more dominant, with ac4C deposition only being significantly restored by secondary A20 and U20 mutations (Fig. 5D, center, Fig. S6B).

Given the apparent anti-correlation between modification and base-pairing, we expected disruption of additional base pairs within U20 variants would enable even more efficient enzyme catalysis. Instead, we observed several mutants in stem I whose effects dominated that of U20 (Fig. 5D, right, Fig. S6C). One striking example came from analysis of position 9, whose mutation was uniformly associated with decreased modification of U20 helices (Fig. 5D, blue lines). In contrast, mutation of its pairing partner 16 increases acetylation (Fig. 5D, orange lines). Since our design embeds *H. sapiens* helix 45 in a larger RNA to facilitate reverse transcription and PCR, we hypothesized co-mutation of U20 and position 9 could be driving formation of a new secondary structure in which helix 45 was disrupted, but the 5’-CCG-3’ substrate was fully base-paired with an upstream 5’-CGG-3’ in the scaffold sequence. In line with this view, in silico structural prediction indicates the A9/U20 mutant is capable of forming an alternative structure in which the 5’-CCG-3’ consensus is housed in a fully paired duplex (‘scaffolded consensus,’ Fig. 5E, Fig. S7). The alternative structure is predicted to be destabilized by mutations in positions 15-17, 23, and 24 in the 3’-stem, potentially explaining the differential effects of mutating the two stems. Further supporting this view, when the modification rate of U20 double mutants is corelated with MFE we find that 5’-stem mutations follow a parabolic distribution, with acetylation of those that highly enforce (C7) or abrogate structure (A9, C9, and U9) disfavored (Fig. S8, blue). This suggests that 5’-stem mutations that are overly destabilizing favor the alternative structure, a relatively poor substrate. In contrast, for 3’-stem mutations MFE and modification are linearly correlated (Fig. S8, green), as would be expected if TkNat10 displayed a straightforward preference for unstructured substrates. Analysis of modification and MFE among all single stem mutants and 3’-stem double mutants (592 data points) further confirms this anti-correlation of modification and negative free energy (Fig. 5F). This preference has some interesting parallels with SHAPE reagents,^25, 26^ reactive chemicals which similarly report on RNA accessibility. Overall, these studies demonstrate the utility of ac4C-seq for high-throughput profiling of RNA acetyltransferase-substrate interactions and specify the 5’-CCG-3’ consensus and RNA secondary structure as important determinants and anti-determinants of in vitro modification by TkNat10.

### Evaluating archaeal enzymes as a tool for sequence-specific RNA acetylation

A major challenge in the study of RNA cytidine acetylation has been a lack of methods to install ac4C into physiological consensus sequences. Recently, we reported a synthetic approach to address this obstacle which enabled the production of sequence-defined RNA oligomers harboring acetylation at specific positions.^27^ One caveat to this approach is that it requires access to custom synthesized phosphoramidite building blocks and significant expertise with automated oligonucleotide synthesis. Considering this, we wondered whether the promiscuous activity of TkNat10 may be amenable to the acetylation of non-native substrates, providing a preparative route to RNAs containing the acetylated 5’-CCG-3’ consensus motif for functional studies. To test this, we incubated a short 10-mer RNA oligonucleotide containing a 5’-CCG-3’ sequence with TkNat10 and cofactors at 65 °C for 3 hours (Fig. 6A). MALDI-TOF confirmed a +42 Da mass shift consistent with to the expected mass of acetylation, which was absent in the enzyme-free control reaction (Fig. 6B).

**Figure 6.**
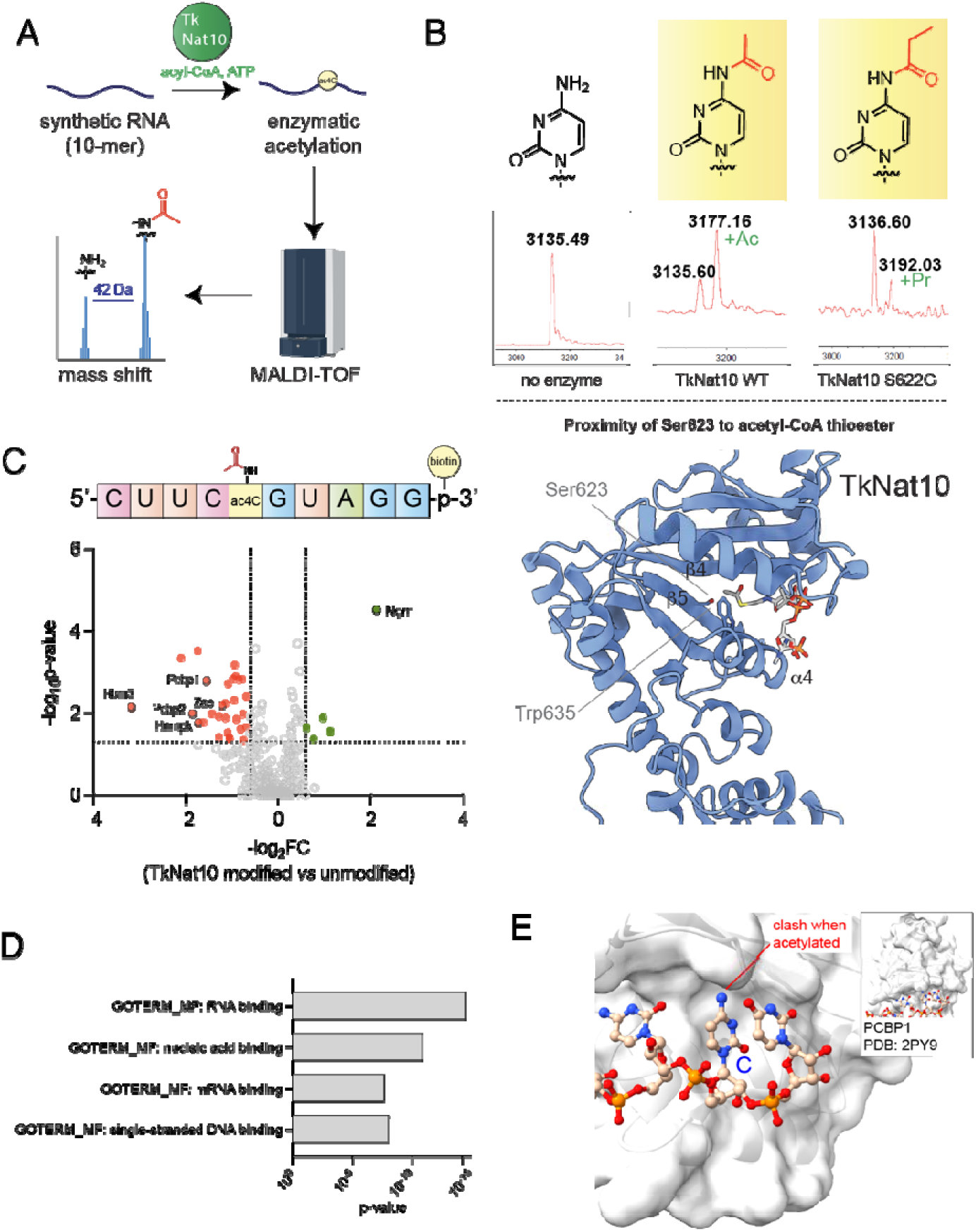
(A) Schematic for MALDI-TOF assay assessing TkNat10 RNA oligonucleotide modification. (B) Top: MALDI-TOF analysis of a single-stranded RNA oligonucleotide treated with no enzyme (left), TkNat10 WT + acetyl-CoA (middle), or TkNat10 S623G + propionyl-CoA (right). Full MALDI traces are provided in the Supplementary Information. Bottom: Predicted structure of TkNat10, indicating bulky residues in active site. Mutation of homologous residues in the human GNAT histone acetyltransferase enzyme KAT2A expands cofactor utilization. (C) LC-MS analysis comparing proteins enriched by a unmodified biotinylated RNA oligonucleotide versus a TkNat10 acetylated RNA oligonucleotide. (D) Gene ontology analysis of proteins preferentially enriched by an unmodified biotinylated RNA oligonucleotide versus a TkNat10-modified acetylated RNA oligonucleotide. (E) Crystal structure of PCBP1 bound to cytidine-containing RNA oligonucleotide. Modeling of N4-acetylation indicates a potential steric clash caused by cytidine modification.

Protein lysine acetyltransferases are able to use alternative acyl-CoA cofactors to catalyze formation of a wide range of acylations including propionylation and crotonylation.^28-30^ The ability of RNA acetyltransferases to catalyze similarly diverse acylations has not been explored. To evaluate TkNat10’s cofactor promiscuity we incubated enzyme with propionyl-CoA and ATP and analyzed modification of our short RNA oligonucleotide substrate by MALDI. Propionyl-CoA was chosen due to its relatively small footprint and known ability by many GNAT-containing protein acetyltransferases.^31^ In contrast to acetyl-CoA, no modification as observed in the propionyl-CoA reaction, suggesting a strict requirement for the endogenous cofactor. To evaluate the molecular basis for this specificity we attempted to relax the specificity by enlarging the active site, expressing and purifying mutants in which Ser623 was mutated to a glycine either alone (S623G) or in combination with Trp635 (S623G/W635A; Fig. 6B, bottom, Fig. S9A). Analogous mutations have been shown to enable the GNAT histone acetyltransferase KAT2A to use biorthogonal azide and alkyne-containing acyl-CoAs.^32^ Analyzing reaction products by MALDI, we found the TkNat10 S623G was able to use propionyl-CoA as a substrate, with the propionylation reaction appearing to proceed to a lesser extent than acetylation (Fig. S9B). Neither enzyme was able to use larger biorthogonal acyl-CoAs as cofactors, and the dual S623G/W635A mutant was completely inactive (Fig. S9C). These results indicate the strict specificity of cytidine acetyltransferases for acetyl-CoA and provide a mutational starting point for using these enzymes to study non-native N4-cytidine acylations.

### Enzymatic modification enables analysis of ac4C-dependent RNA-protein interactions

Nucleobase modifications can act as potent regulators of RNA-protein interactions.^33, 34 35^ Previous studies have demonstrated that using in vitro transcription to homogenously replace cytidine with N4-acetylcytidine in synthetic mRNAs can alter the RNA-protein interactomes of these molecules.^36^ However, this approach presents several caveats. It homogenously installs ac4C in a variety of non-native contexts, cannot access a 5’- C**<u>C</u>**G-3’ sequence in which ac4C is flanked by an unmodified cytidine nucleotide, and produces a variety of byproducts such as double-stranded antisense transcripts that may confound results.^37^ To overcome these limitations, studies of RNA modifications such as N1-methylpseudouridine (m1Y) have benefitted from the use of synthetic m1Y-containing oligonucleotides to determine, for example, how this modification effects the immune response.^38^ Since short synthetic ac4C-containing oligonucleotides are not available to most researchers, we asked: could we use post-synthetic enzymatic modification to study how ac4C affects protein-RNA interactions?

To address this, a biotinylated 10-mer RNA containing a 5’-CCG-3’ sequence was obtained via commercial phosphoramidite synthesis. This substrate was incubated with TkNat10 and cofactors at 65 °C for 3 h and acetylation was verified by MALDI (Fig. S10). To test ac4C’s ability to affect recognition by RNA-binding proteins (RBPs), we individually incubated two biotinylated RNAs – one with TkNat10-catalyzed ac4C and one without - with RAW 264.7 cell lysates, enriched over streptavidin, and analyzed the proteins captured by LC-MS/MS (Fig 6C, Table S3). Focusing on proteins identified by >1 peptide, we identified 18 proteins whose capture was significantly impeded by ac4C (p-value ≤ 0.05, Log2 fold change ≤ -1) and 4 proteins whose capture was facilitated by ac4C. Interestingly, the majority of proteins whose capture was impeded by ac4C were RBPs (Fig 6D), while the majority of proteins whose capture was facilitated by ac4C have been identified as common contaminants in proteomics experiments.^39^ One potential explanation for the latter observation is that RNA acetylation impedes RBP/resin interactions, which are then saturated by non-specific protein binding. Among the RBPs whose interaction is disrupted by ac4C are known PCBP1, PCBP2, and ZC3HAV1, all of which have demonstrated evidence of cytidine-dependent RNA-protein interactions.^40, 41^ The potential for ac4C to interfere cytidine-dependent protein binding is illustrated by modeling the modification into a crystal structure of RNA bound to PCBP1 (Fig 6E). Our findings provide further evidence for the ability of cytidine acetylation to influence RNA interactomes and highlight archaeal cytidine acetyltransferase enzymes as a powerful tool for functional analysis of 5’-CCG-3’ acetylation.

## Discussion

Here we report the reconstitution, biochemical profiling, and functional application of a *T. kodakarensis* RNA acetyltransferase. Our studies shed light on unanswered questions around these catalysts, including the specific relevance of their acetyltransferase activity to archaeal fitness, their substrate and cofactor selectivity, and potential applications in preparative production of ac4C oligonucleotides. We demonstrate the utility of quantitative ac4C sequencing for high-throughput profiling of RNA acetyltransferase substrates. This study confirmed TkNat10’s preference for single-stranded RNA substrates, suggesting a potential application of this enzyme as a structural probe. Finally, we show this promiscuous catalyst can be applied to N4-acetylate the 5’-CCG-3’ consensus motif short RNA oligonucleotide contexts and use this to examine the impact of site-specific ac4C on RNA-protein interactions.

Our results also highlight features warranting further investigation. One surprising result was the apparent incongruity between the preferred cellular versus biochemical targets of TkNat10. Applying an identical ac4C sequencing method, we found that in cells rRNA and tRNAs are the most highly modified targets of the enzyme,^7^ while in biochemical experiments unstructured RNA substrates are preferred. Several factors may contribute to this observation. First, our assay conditions undoubtedly present different concentrations of cofactors and ions than the endogenous intracellular environment of *T. kodakarensis*. Along similar lines, our recombinant enzyme preparations were applied to a deproteinized *T. kodakarensis* transcriptome, which is expected to have a large impact on the presentation of rRNA substrates. Perhaps the most intriguing hypothesis for this dissimilarity may come from consideration of eukaryotic RNA acetylation, where short nucleolar RNA (snoRNA) and protein adapters are used to guide ac4C to specific sites.^11, 12^ Archaea have equivalent sno-like RNA (sRNA) mechanisms which help carry out modifications such as 2’-O-methylation.^42^ Future studies aiming to define the input of sRNAs as well as other proteins into archaeal ac4C deposition should help to better understand this phenomenon. TkNat10’s preference for unfolded substrates is also interesting in light of recent studies indicating ac4C can serve to enforce RNA secondary structure.^7, 27^ This could conceivably allow the stand-alone enzyme to perform a sentinel function by sensing unfolded RNAs and installing a modification to promote folding and – potentially - fitness. Testing this hypothesis will benefit from additional research exploring ac4C’s capacity to function as an RNA structure switch, which should in turn be facilitated by the enzymatic modification strategies described here.

It is also important to specify some limitations of our study and future work that may help address them. Our high-throughput studies explored TkNat10’s activity within the context of only two hairpin substrates and used MFE as a simple proxy for RNA structure. Integrating analysis of more diverse substrates structure probing data could provide a more comprehensive understanding of archaeal RNA acetyltransferase activity and guide future applications. A barrier faced in our biochemical characterization was the requirement for heating, which hindered application of conventional coupled enzyme techniques to determine kinetic constants for TkNat10. We anticipate this may be remedied in the future by using stop-point assays in combination with highly specific and sensitive tools for acetyl-CoA or CoA detection.^43, 44^ Finally, while we were able to demonstrate near stoichiometric enzymatic modification of a short RNA oligonucleotide, TkNat10’s utility as a tool for site-specific RNA acetylation is currently limited to contexts where a consensus motif exists. Efforts to broaden the substrate selectivity of this enzyme may benefit from directed evolution strategies capable of converting modifications into functional changes in RNA or protein activity, examples of which have been reported.^45^ Overall, our studies provide new insights into the biochemistry of a highly conserved enzyme family and a foundation for studying the role of site-specific RNA acetylation in biology, therapeutics, and disease.

## Data availability

Custom perl scripts used in this study can be found at the GitHub repository, https://github.com/Meier-Lab-NCI/TkNat10.

## Acknowledgements

We thank Ronald Holewinski and Thorkell Andresson (Protein Characterization Laboratory, Frederick National Laboratory for Cancer Research) for LC/MS/MS and proteomic analysis. This work is supported by the Intramural Research Program of the National Institutes of Health (NIH), the National Cancer Institute, The Center for Cancer Research (ZIA BC011488) (J.L.M.). S. S. is funded by the Israel Science Foundation (grant no. 913/21), the European Research Council (ERC) under the European Union’s Horizon 2020 research and innovation programme (grant no. 101000970). A.S.C. is supported by the Israel Science Foundation (grant no. 1260/23) and by the Azrieli Foundation. T.J.S is supported by the USA National Science Foundation, award #2022065. L. A. was supported by the Intramural Research Program of the National Library of Medicine at the National Institutes of Health. In addition, this research was carried out in part with federal funds from the National Cancer Institute, National Institutes of Health, under contract HHSN261200800001E (J. K.). The content of this publication does not necessarily reflect the views or policies of the Department of Health and Human Services, nor does mention of trade names, commercial products or organizations imply endorsement by the US Government.

## Supporting Information

**Figure S1.**
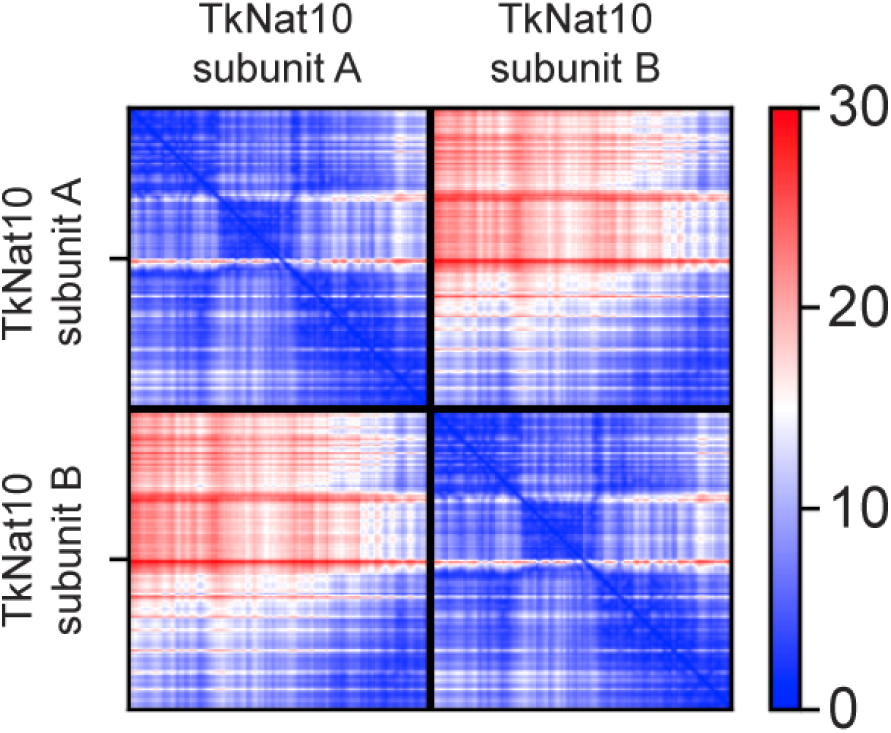
AF-Multimer-generated 2D plot of predicted alignment error from predictions of TkNat10 dimeric interaction. Predicted interactions are found in the top right and bottom left panels, where blue corresponds to lower estimated error suggestive of interaction.

**Figure S2.**
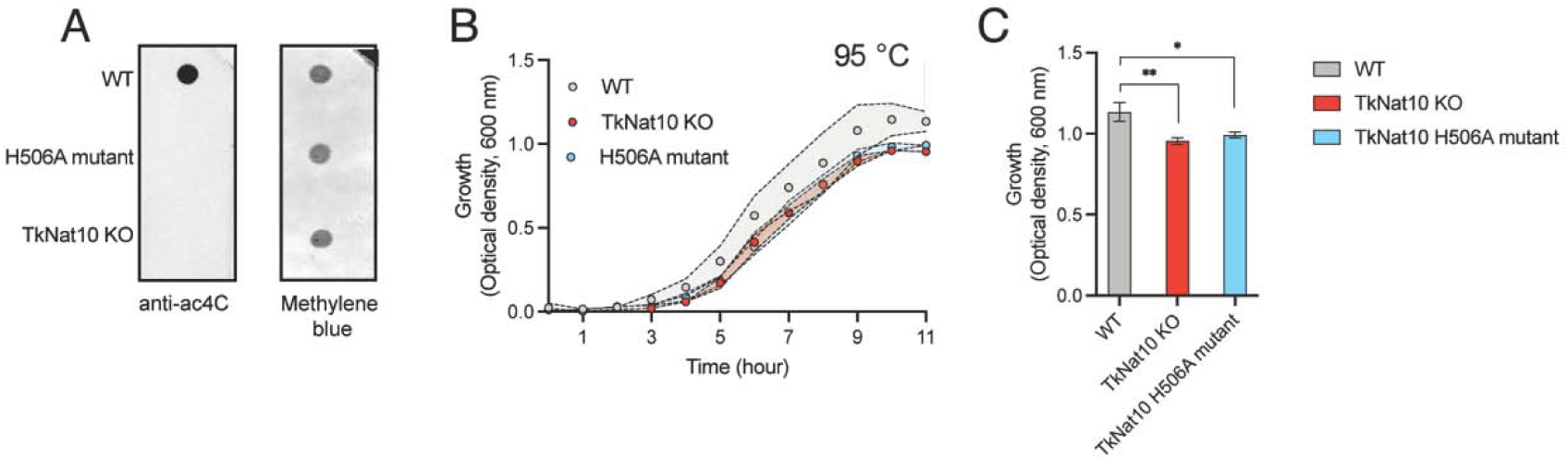
(A) Dot blot confirming loss of ac4C in TkNat10 catalytic H506A mutant at 85 °C. (B) Growth rate of WT *T. kodakarensis*, TkNat10 deletion strain, and strain containing a TkNat10 H506A catalytic mutant at 95 °C. (C) Differential growth of *T. kodakarensis* strains at 95 °C 11 hours.

**Figure S3.**
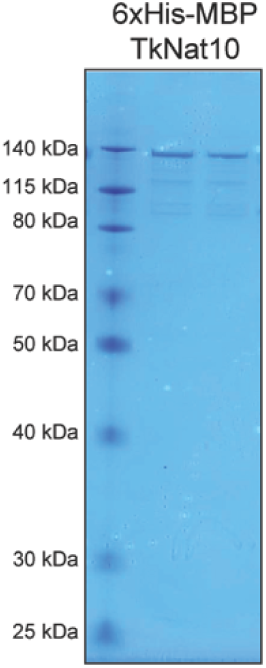
SDS-PAGE gel of purified 6xHis-MBPTkNat10 enzyme.

**Figure S4.**
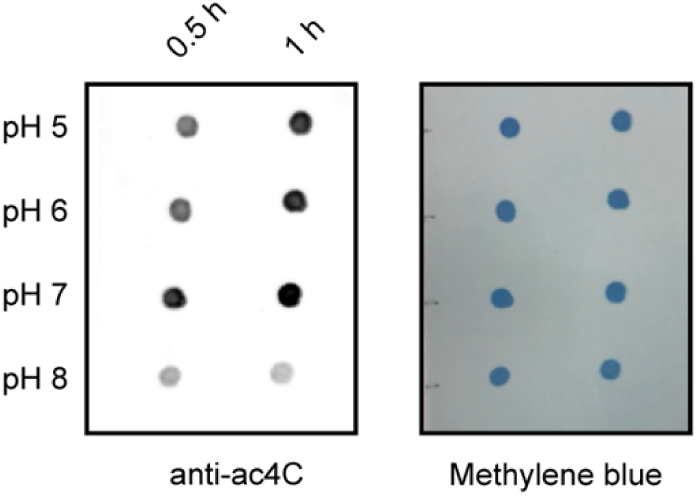
Dependence of recombinant TkNat10 activity on pH.

**Figure S5.**
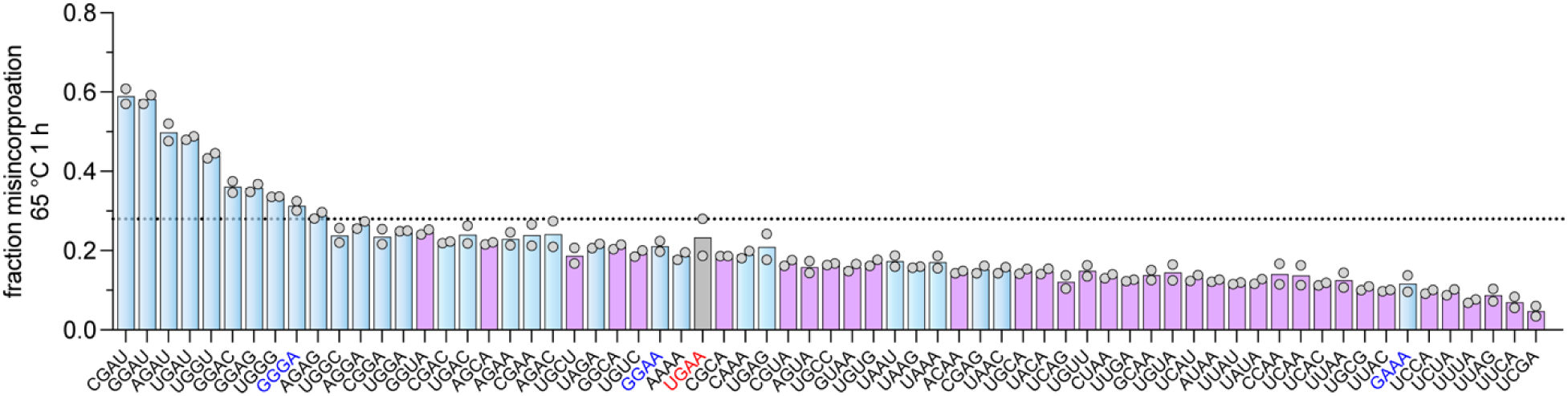
Fraction misincorporation observed upon incubation of TkNat10 and cofactors with 18S rRNA variants containing a native 5’ and 3’ stem in combination with the specified tetraloop sequences (65 °C, 1 h). The native 5’-UGAA-3’ tetraloop sequence is specified in red, stable 5’-GNRA-3’ tetraloops in dark blue, tetraloops with internal purines (5’-NRRN-3’) in light blue and all other tetraloops are in purple. The dashed line corresponds to the highest fraction misincorporation value observed for the native 5’-UGAA-3’ tetraloop.

**Figure S6.**
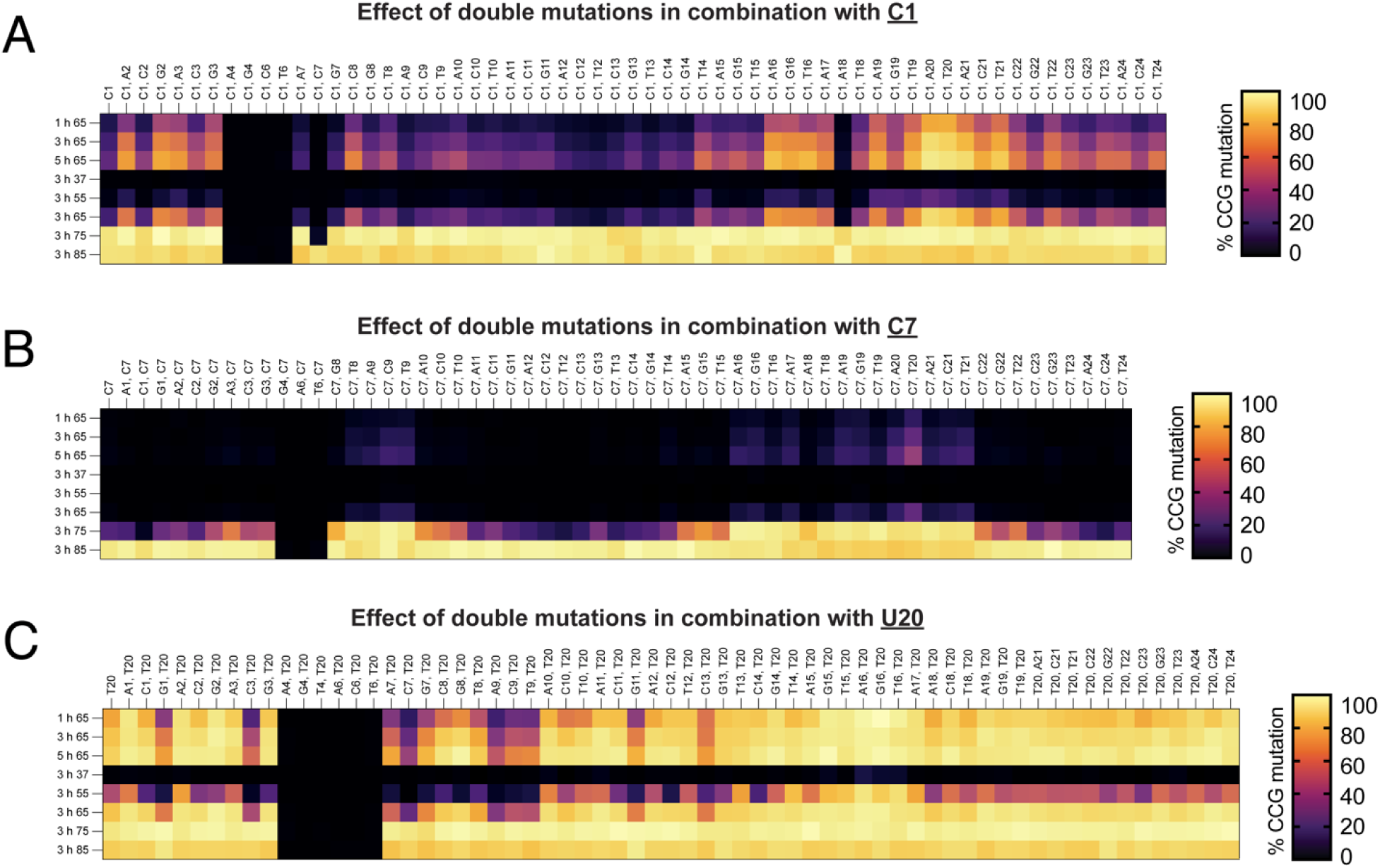
(A) Heatmap depicting % misincorporation (‘% CCG mutation’) as a function of time and temperature for 18S rRNA substrates with a C1 mutation. Each column corresponds to a mutant sequence with the mutation indicated in the top row of the header and the position in the bottom row of the header. (B) Equivalent data for 18S rRNA substrates with a C7 mutation. (C) Equivalent data for 18S rRNA substrates with a U20 mutation. Data represents the average of two replicate experiments. Raw data is provided in Table S2.

**Figure S7.**
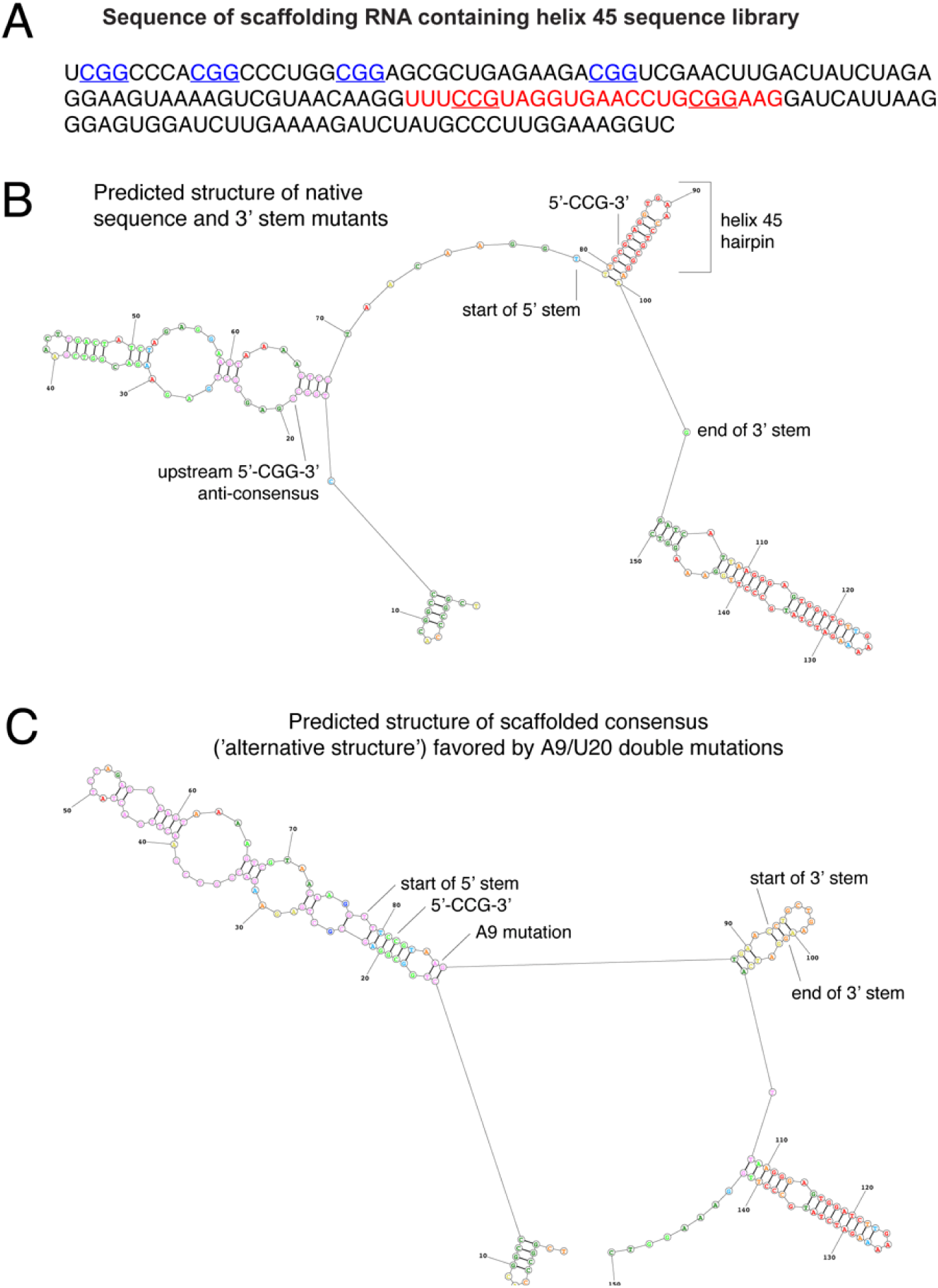
(A) Description of scaffolding sequence flanking helix 45 rRNA libraries. (B) Predicted structure of native helix 45 substrate. (C) Predicted structure of A9/U20 double mutant. Structures were analyzed using the RNAstructure web server with default settings, which can be accessed at: https://rna.urmc.rochester.edu/RNAstructureWeb/Servers/Predict1/Predict1.html

**Figure S8.**
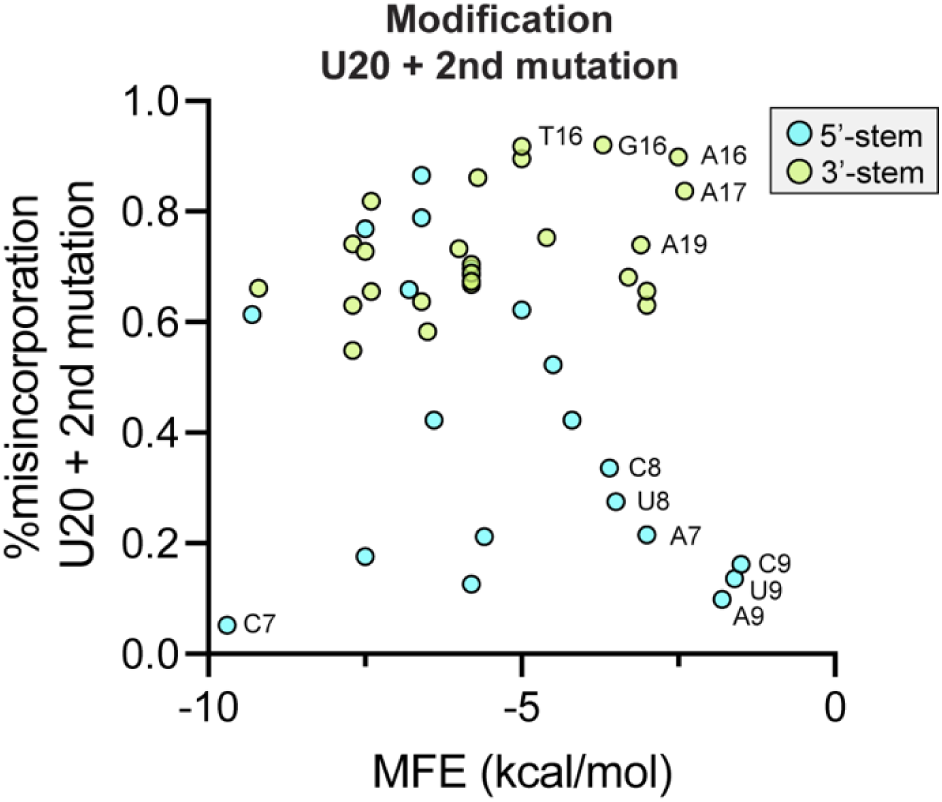
(A) Correlation of ac4C-seq misincorporation percentage with minimum free energy (MFE) for 18S rRNA U20 variants. U20 double mutants containing secondary mutations in the 5’ stem display a parabolic modification profile (blue), with acetylation of unstructured substrates presumably disfavored due to rearrangement into an alternative structure in which the 5’-CCG-3’ substrate is fully paired with a sequence in the upstream scaffold. U20 double mutants containing secondary mutations in the 3’ stem show a linear correlation (green) with MFE as would be expected if TkNat10 preferred unstructured substrates.

**Figure S9.**
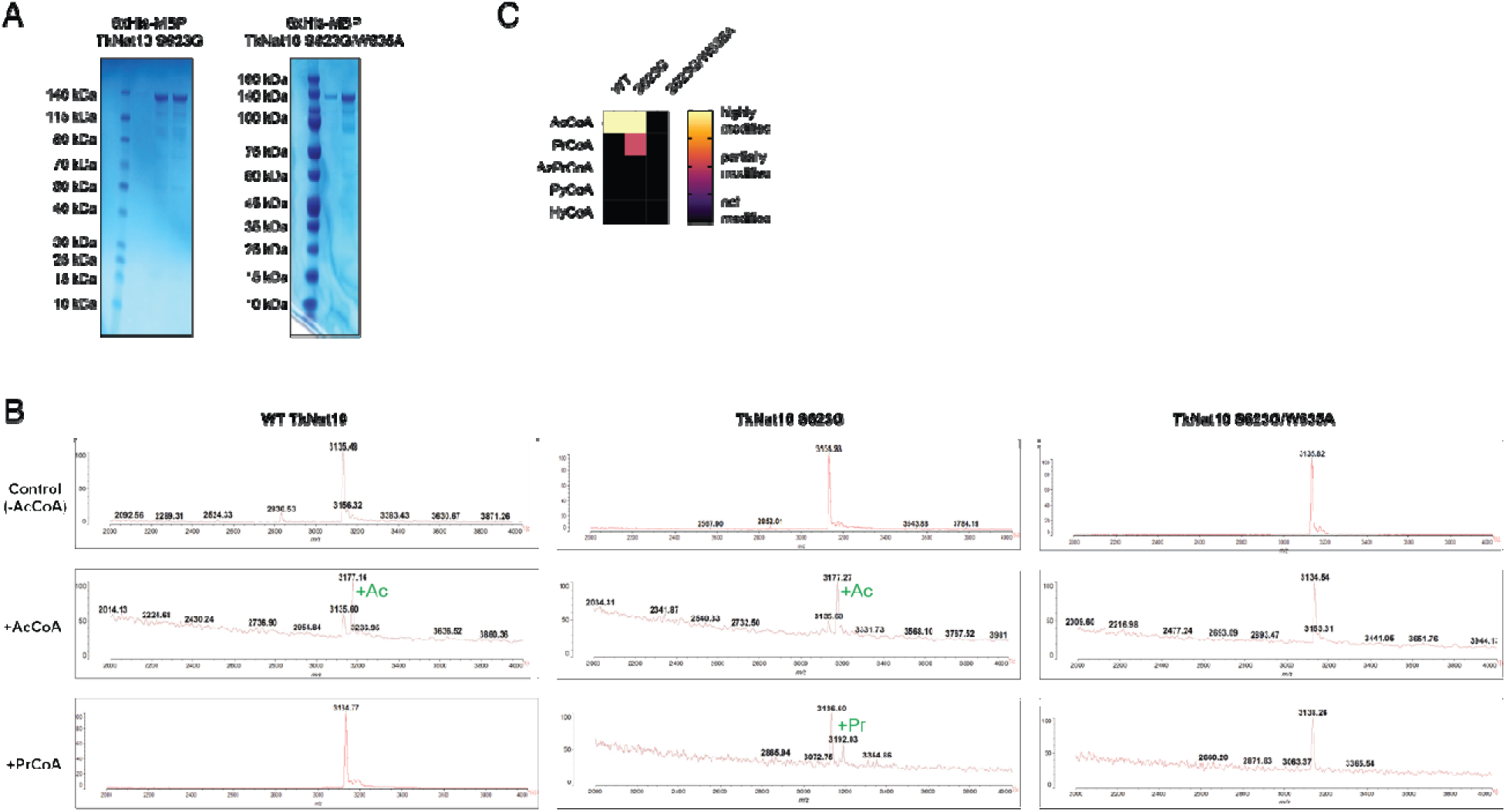
(A) SDS-PAGE gels of TkNat10 S623G and S623G/W635G double mutants. (B) Complete MALDI-TOF mass spectra of single-stranded RNA oligonucleotide treated with WT TkNat10, S623G, and S623G/W635A mutants in the presence of Acetyl CoA (middle) and propionyl CoA (bottom). Control reactions without CoA cofactors are shown for comparison (Top). (C) Heat map depicting activity of native TkNat10 and mutants with alternative acyl-CoA cofactors.

**Figure S10.**
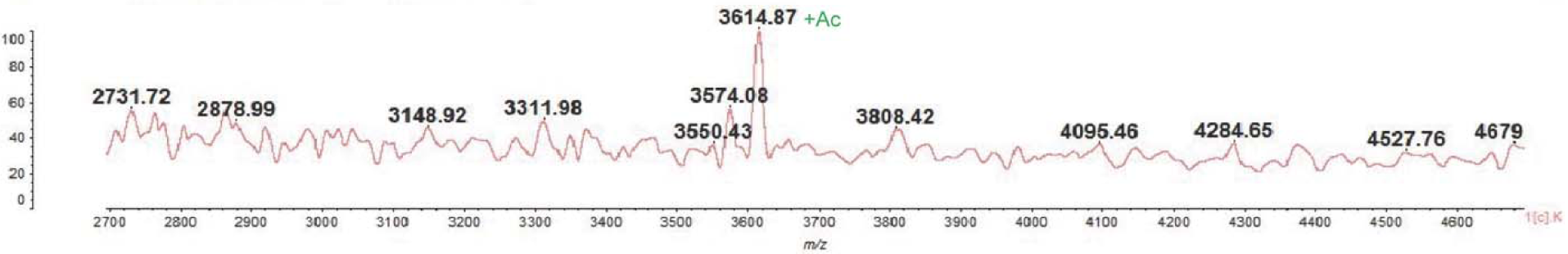
The complete MALDI-TOF mass spectrum of the acetylated RNA-Biotin (m/z: 3614.87) used in the RNA-protein pull-down experiment. Unreacted RNA-Biotin 10-mer appears at m/z=3574.08.

